# Standing genetic variation of the AvrPm17 avirulence gene in powdery mildew limits the effectiveness of an introgressed rye resistance gene in wheat

**DOI:** 10.1101/2021.03.09.433749

**Authors:** Marion C. Mueller, Lukas Kunz, Seraina Schudel, Sandrine Kammerecker, Jonatan Isaksson, Michele Wyler, Johannes Graf, Alexandros G. Sotiropoulos, Coraline R. Praz, Thomas Wicker, Salim Bourras, Beat Keller

**Affiliations:** Department of Plant and Microbial Biology, University of Zurich, Zurich, Switzerland; Department of Forest Mycology and Plant Pathology, Division of Plant Pathology, Swedish University of Agricultural Sciences, Uppsala, Sweden

**Author notes:** Authors contributed equally (Marion C. Mueller, Lukas Kunz). Corresponding authors: Beat Keller, Salim Bourras **Email**. **Author Contributions** M.C.M., L.K., S.B. and B.K. wrote the paper. S.B., and B.K. coordinated the research. M.C.M., L.K.,S.B. and B.K. designed the research. M.C.M. and S.S. performed genetic analysis. M.C.M., L.K.,S.S. S.K. J.I. performed experiments. M.C.M, S.S., M.W., C.R.P. A.G.S, J.G. and T.W. performed bioinformatics analysis.

**Keywords:** wheat, powdery mildew, resistance introgression, gene conversion, avirulence gene

## Abstract

Introgressions of chromosomal segments from related species into wheat are important sources of resistance against fungal diseases. The durability and effectiveness of introgressed resistance genes upon agricultural deployment is highly variable - a phenomenon that remains poorly understood as the corresponding fungal avirulence genes are largely unknown. Until its breakdown, the *Pm17* resistance gene introgressed from rye to wheat provided broad resistance against powdery mildew (*Blumeria graminis*). Here, we used QTL mapping to identify the corresponding wheat mildew avirulence effector *AvrPm17*. It is encoded by two paralogous genes that exhibit signatures of re-occurring gene conversion events and are members of a mildew sub-lineage specific effector cluster. Extensive haplovariant mining in wheat mildew and related sub-lineages identified several ancient virulent *AvrPm17* variants that were present as standing genetic variation in wheat powdery mildew prior to the *Pm17* introgression, thereby paving the way for the rapid breakdown of the *Pm17* resistance. QTL mapping in mildew identified a second genetic component likely corresponding to an additional resistance gene present on the 1AL.1RS translocation carrying *Pm17*. This gene remained previously undetected due to suppressed recombination within the introgressed rye chromosomal segment. We conclude that the initial effectiveness of 1AL.1RS was based on simultaneous introgression of two genetically linked resistance genes. Our results demonstrate the relevance of pathogen-based genetic approaches to disentangle complex resistance loci in wheat. We propose that identification and monitoring of avirulence gene diversity in pathogen populations becomes an integral part of introgression breeding to ensure effective and durable resistance in wheat.

**Significance Statement:** Domesticated and wild wheat relatives provide an important source of new immune receptors for wheat resistance breeding against fungal pathogens. The durability of these resistance genes is variable and difficult to predict, yet it is crucial for effective resistance breeding. We identified a fungal effector protein recognised by an immune receptor introgressed from rye to wheat. We found that variants of the effector allowing the fungus to overcome the resistance are ancient. They were already present in the wheat powdery mildew gene pool before the introgression of the immune receptor and are therefore responsible for the rapid resistance breakdown. Our study demonstrates that the effort to identify new resistance genes should be accompanied by studies of avirulence genes on the pathogen side.

## Main Text

### Introduction

Wheat is the most widely cultivated food crop and is susceptible to a number of fungal diseases. For more than a century, breeding for genetically resistant cultivars that can durably withstand disease has been one of the most important approaches for sustainable wheat production globally. Introgressions of chromosomal segments from closely related wild grasses such as *Aegilops* or *Agropyron* species (1, 2) and other crop species such as rye (*Secale cereale*) have been highly valuable sources of new resistance gene specificities (3). Specifically, the 1BL.1RS or 1AL.1RS translocations of the rye chromosome 1R introgressed into hexaploid (AABBDD) wheat (reviewed in (4)) were of great relevance for wheat resistance breeding. Genes present on these translocations are widely used in wheat breeding and confer resistance to leaf rust (*Lr26*), stripe rust (*Yr9*), stem rust (*Sr31, Sr50/SrR, Sr1R*^*Amigo*^) and powdery mildew (the allelic *Pm8*/*Pm17* pair), (5, 6).

It has been proposed that introgressed resistance genes provide more effective and potentially more durable resistance since pathogens specialized on wheat have previously not been exposed and therefore have not adapted to the resistance specificities that evolved in other species (7). This is exemplified by the rye *Sr31* gene, which was deployed worldwide. It provided effective and broad resistance against *Puccinia graminis* f. sp. *tritici* the causal agent of wheat stem rust for over 30 years before being overcome by the virulent African strain Ug99 (8), demonstrating both the huge benefit of an introgressed rye gene as well as the constant need for new broadly active resistance genes (9). In contrast to *Sr31* and the general hypothesis, many introgressed resistance genes were overcome quickly by wheat pathogens (10). For example, the rye introgressions with *Pm8* and *Pm17* became ineffective against wheat powdery mildew *Blumeria graminis f. sp. tritici* (*B*.*g. tritici*) within a few years after their deployment in large-scale agricultural settings (11-14). Thus, it remains one of the most pressing questions in the field of plant breeding research why a few introgressed genes such as *Sr31* remained effective over a long timeframe and despite worldwide deployment whereas others are overcome quickly (7).

The allelic *Pm8* and *Pm17* genes encode for nucleotide-binding leucine rich repeat (NLR) proteins that were introgressed into wheat from ‘Petkus’ and ‘Insave’ rye cultivars respectively (15, 16). It was demonstrated that both genes represent rye homologs of the wheat *Pm3* resistance gene (15, 16) which encodes for a high number of different NLR alleles that confer race-specific resistance against wheat powdery mildew through recognition of mildew encoded avirulence proteins (17-19).

In wheat powdery mildew recent studies using map-based cloning, GWAS and effector benchmarking approaches have identified several avirulence genes, among them *AvrPm3*^*a2/f2*^, *AvrPm3*^*b2/c2*^ and *AvrPm3*^*d3*^ recognized by *Pm3a/Pm3f, Pm3b/Pm3c* and *Pm3d* respectively (17, 18). Sequence analysis of wheat mildew *Avr* genes revealed that they all encode small secreted candidate effector proteins (17, 18, 20, 21) and exhibit high levels of sequence variation on a population level including the independent evolution of numerous gain of virulence alleles by diverse molecular mechanisms (17, 18, 20, 22). The identification and functional characterization of mildew avirulence genes has therefore significantly broadened our understanding of race-specific resistance and resistance gene breakdown in the wheat – mildew pathosystem.

Grass powdery mildews exist in many sub-lineages also called *formae speciales* (*f*.*sp*.) that are highly host specific such as mildew on wheat (*B*.*g. tritici*), rye (*B. g. secalis*) or the wheat/rye hybrid triticale (*B*.*g. triticale*) which emerged recently and was attributed to a hybridization event between wheat and rye mildew sub-lineages (23, 24). Due to the strict host barrier, it is assumed that non-adapted mildew sub-lineages have not been exposed to NLR resistance specificities of an incompatible host and therefore have not evolved to evade recognition. Indeed, several *Pm3* alleles have been found to contribute to non-host resistance through recognition of conserved avirulence effectors in non-adapted mildew sub-lineages such as *B*.*g. secalis* (18). Given these observations, the rapid breakdown of *Pm8* and *Pm17* resistance after introgression into wheat remains puzzling and provides an opportunity to study evolutionary dynamics of wheat mildew in the context of introgression breeding. The *Pm17* introgression is especially suited for this purpose since the associated 1AL.1RS translocation, first described in 1976 in Oklahoma (US) (25), was not used before the end of the 20^th^ and has been deployed in large-scale agricultural setting only in the beginning of the 21^th^ century in the United States, where it provided resistance against wheat mildew in bread wheat (26, 27). In contrast, the deployment in other wheat growing areas globally started only after the year 2000 (11, 27-29), and breakdown of *Pm17* resistance was generally observed within few years and has been well documented in several wheat growing regions such as the US, China and Switzerland (11, 13, 27, 30).

In this study we report the molecular basis underlying the resistance breakdown of the introgressed *Pm17* gene in wheat. Using QTL mapping in a bi-parental mildew population we demonstrate that the corresponding avirulence effector *AvrPm17* is encoded by a paralogous effector gene pair, residing in a dynamic effector cluster, specific to the wheat and rye mildew sub-lineages. Moreover, we describe the identification of numerous ancient virulence alleles of the *AvrPm17* gene that have been present as standing genetic variation in *B*.*g. tritici* even before the introgression of *Pm17* into the wheat breeding pool. Lastly, we provide genetic evidence for the existence of a so far unidentified resistance gene against wheat mildew which was co-introgressed with *Pm17* from rye, which could be revealed through careful dissection of resistance specificities based on genetic studies in the pathogen.

## Results

### QTL mapping identifies a single avirulence locus for Pm17 in wheat powdery mildew

To understand the breakdown of the rye NLR *Pm17* in wheat we aimed at identifying its corresponding avirulence gene by taking advantage of the recent cloning of *Pm17* and its validation in transgenic wheat lines (16). We used a preexisting, sequenced F1 mapping population derived from a cross of the avirulent *B*.*g. triticale* isolate THUN-12 and *B*.*g. tritici* isolate Bgt_96224 which exhibits a virulent phenotype on the independent transgenic lines Pm17#34 and Pm17#181 (Fig. S1A,B), (31). A single interval QTL mapping approach using 55 randomly selected progeny of the Bgt_96224 X THUN-12 cross identified a single locus on chromosome 1 at identical map position (164.8cM) with highly significant LOD scores of 9.2 for wheat genotype Pm17#34 and 7.0 for Pm17#181 respectively (Fig. 1A-C, Fig S1C,D, Table S1). The pericentromeric location of the mapped *AvrPm17* locus contrasts with the location of previously identified wheat mildew avirulence genes that tend to reside near the telomeric region or on the chromosome arms (Fig. 1B, Fig. S2, (17, 18, 20)).

**Figure 1.**
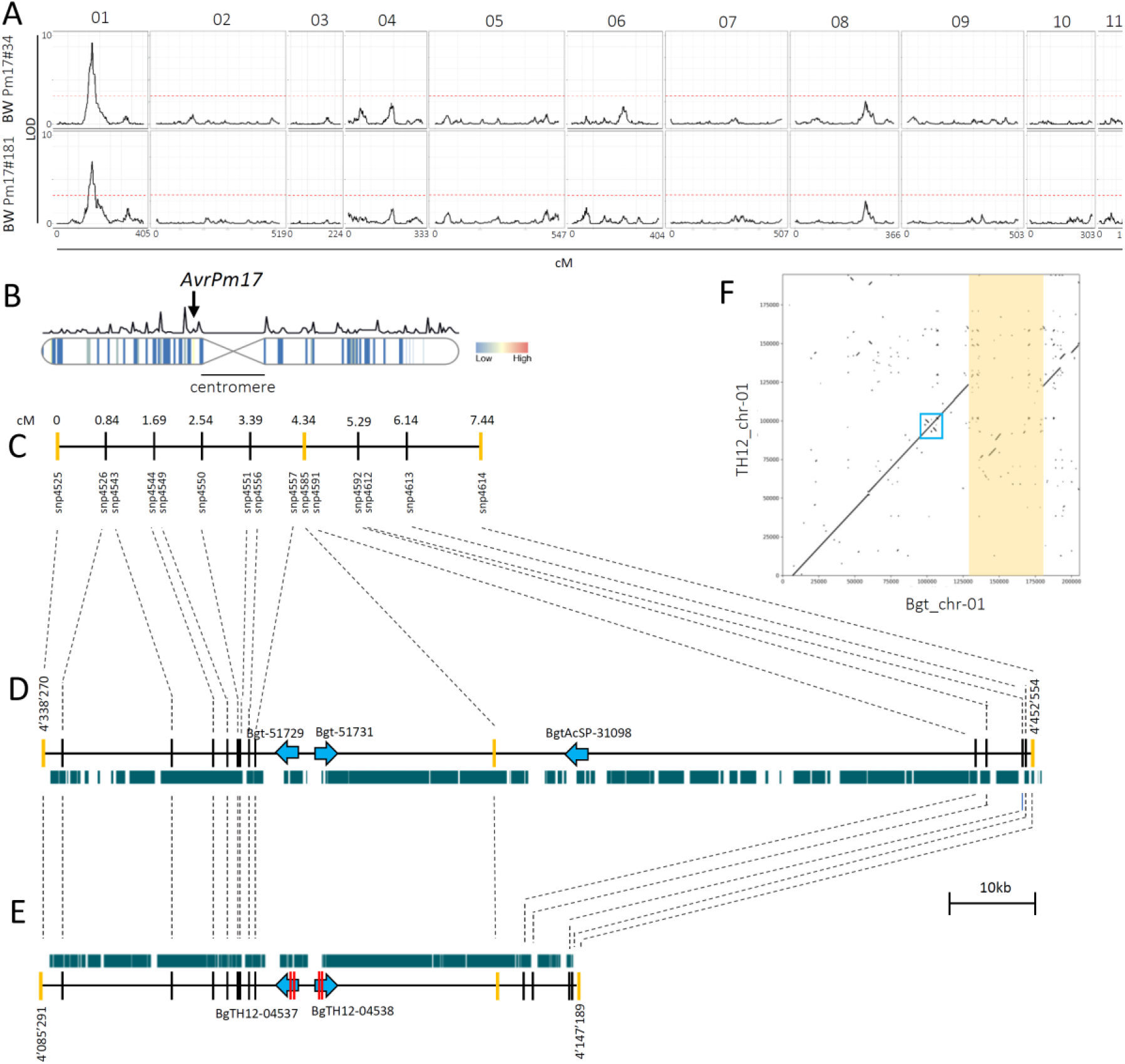
Avirulence on wheat genotypes with *Pm17* is controlled by a single locus in a biparental mapping population between *B*.*g. tritici* 96224 X *B*.*g. triticale* THUN-12. (A) Single interval QTL mapping of 55 progeny of the cross 96224X THUN-12 on two transgenic lines expressing *Pm17*-HA under control of the maize ubiquitin promoter (Ubi) promoter. The genetic map of Bgt_96224 X THUN-12 based on 119,023 markers was published previously in (31) and contains eleven linkage groups that correspond to the eleven chromosomes of *B*.*g. tritici* 96224 and *B*.*g. triticale* THUN-12 ((31), unpublished data) Significance level of the LOD (logarithm of the odds) value was determined using 1000 permutations and is indicated by a red line. (B) Location of the QTLs identified in the pericentromeric region of chromosome 1 of *B*.*g. tritici* 96224. The centromeric region is indicated. Vertical bars indicate effector gene density in 50kb windows following a gradient indicated in the color key.. Line above the chromosome indicates the recombination rate in cM/50kb as described in (31). The remaining ten chromosomes are depicted in Fig. S2. (C) Informative markers in the 7.44 cM genetic confidence interval (1.5LOD) and their cM position relative to the left flanking marker. Flanking markers and the best associated marker of the QTL are depicted in yellow. (D-E) Physical interval underlying the genetic interval in *B*.*g. tritici* 96224 (D) and *B*.*g. triticale* THUN12 (E). Gene and gene orientation are indicated with blue arrows (gene length not drawn to scale). Non-synonymous SNPs in THUN-12 versus isolate 96224 are indicated by a red bar within the gene. (D-E) Green bars indicates the presence of transposable elements in the interval. (F) Alignment showing the region flanking 1Mb up and downstream of the *AvrPm17* locus in the reference assemblies of *B*.*g. tritici* 96224 and *B*.*g. triticale* THUN-12. The location of the paralogous effectors *BgTH12-04537*/*BgTH12-04538* and *Bgt-51729/Bgt-51731* is indicated by a blue box. The 50kb deletion in THUN-12 compared to 96224 is highlighted in yellow.

To identify *AvrPm17* candidate genes we analysed the physical region underlying the QTL (with a confidence interval of 1.5 LOD) on chromosome 1 in the chromosome-scale assemblies of the parental isolates Bgt_96224 and THUN-12 ((31), unpublished data). The genetic confidence interval corresponded to a 61.8kb region in the assembly of THUN-12 and a much larger region of 114.3 kb in the assembly of Bgt_96224. This striking difference in size is explained by a large 50kb deletion in the THUN-12 genome (Fig. 1D-F). The interval in the *Pm17* avirulent isolate THUN-12 encodes only encodes a paralogous effector gene pair *BgTH12-04537* and *BgTH12-04538* (Fig. 1E). The two effector genes encode for identical proteins that differ by two synonymous single nucleotide polymorphisms (SNPs). The two gene copies are encoded by two inverted duplicated segments of 2’300bp separated by a 4’769bp intergenic region (Fig. 1F). The corresponding gene duplication is also present in the corresponding region of the virulent parent Bgt_96224. There, the duplicated effector genes *Bgt-51729* and *Bgt-51731* are identical and each carries two amino acid changes (A53V, R80S) compared to BgTH12-04537 and BgTH12-04538 respectively (Fig. 1D-E, Fig. 2A, Fig. S3). The interval in the Bgt_96624 genome encodes an additional effector gene; *BgtAcSP-31098* that lies within the 50kb deleted region in THUN-12. In the absence of additional genes in the locus of the avirulent parent THUN-12 we predicted that *BgTH12-04537* and *BgTH12-04538* encode for *AvrPm17*. Using RNA-seq data from the parental isolates we found that *BgTH12-04537/BgTH12-04538* and *Bgt-51729/Bgt-51731* are highly expressed at early stages of infection corresponding to the establishment of the haustorial feeding structure at 2dpi, reminiscent of other wheat mildew *Avr* genes (Fig. S4 and S5). The *AvrPm17* candidates are not differentially expressed (logFC<1.5, Fig. S4) therefore indicating that the amino acid polymorphisms observed between Bgt_96224 and THUN-12 must account for the difference in phenotype.

**Figure 2.**
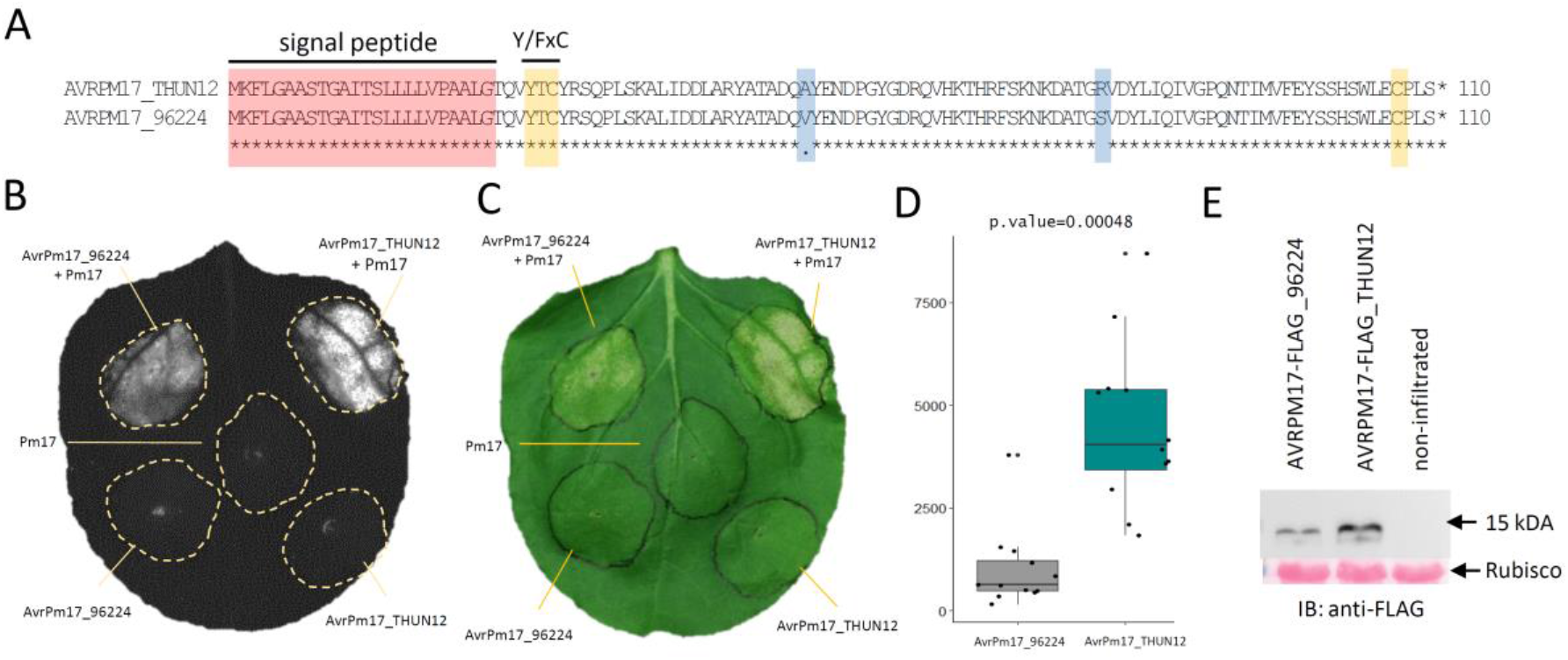
Functional validation of *AvrPm17* in *N. benthamiana* (A) Protein alignment of the AVRPM17 candidate in THUN-12 and 96224. Predicted signal peptide, Y/FxC motif and C-terminal cysteine residues are indicated in red and yellow, respectively. Polymorphic amino acid residues between Bgt_96224 and THUN-12 are highlighted in blue. (B-C) Co-expression of *Pm17*-HA and *AvrPm17*_THUN12 (*BgTH12-04537/BgTH12-04538*) and *AvrPm17*_96224 (*Bgt-51729/Bgt-51731*) by transient Agrobacterium-mediated expression in *Nicotiana benthamiana*, imaged by the Fusion FX imager system (B) or a conventional camera (C). Co-infiltrations were done at a ratio of 1:4 R:Avr. (D) Difference in hypersensitive response induction between the two AVRPM17 variants AVRPM17_THUN12 and AVRPM17_96224 infiltrated in a ratio of R:Avr of 1:1. The p-values of the paired Wilcoxon-ranked sum test is indicated above the panel. (E) Western blot showing C-terminal tagged AVRPM17-FLAG variants extracted from *A. tumefaciens* infiltrated leaf areas of *N. benthamiana* at 2dpi.

A previous study found that the hybrid genome of *B*.*g. triticale* isolates consists of distinct genomic segments inherited from either wheat or rye mildew (23). Due to the rye origin of the *Pm17* resistance gene, the origin of the avirulence locus in triticale mildew THUN-12 is of special interest. Following the approach of (23) based on the analysis of fixed polymorphism between wheat and rye mildew, we found that the physical region underlying the *AvrPm17* QTL in THUN-12 is a segment inherited from wheat mildew (Fig. S6A-C). This indicates that that the rye *Pm17* gene recognises an avirulence component originating from the non-adapted wheat powdery mildew donor and not from the adapted rye mildew, in the triticale mildew hybrid.

### Functional validation of *AvrPm17*

To functionally validate *AvrPm17*, we transiently co-expressed the *BgTH12-04537/BgTH12-04537* Avr candidate with *Pm17*-HA in *Nicotiana benthamiana* by *Agrobacterium tumefaciens* mediated transient overexpression (18, 20). All effector constructs were expressed without the signal peptide and codon-optimized for expression in *N. benthamiana* to ensure optimal translation *in planta. BgTH12-04537/BgTH12-04538* elicited a strong hypersensitive response (HR) upon co-expression with Pm17-HA but not when expressed alone, confirming that these paralogous effector genes are *AvrPm17* (Fig. 2B,C). Co-expression of *AvrPm17_THUN12* with either the *Pm8* gene from rye or the *Pm17* orthologues from wheat (*Pm3a-f, Pm3CS*) did not result in a hypersensitive response in *N. benthamiana* (Fig. S7), demonstrating the specificity of *AvrPm17_THUN-12* recognition by *Pm17*.

Interestingly-co-expression of *AvrPm17_96224* with *Pm17*, also resulted in a hypersensitive response. The extent of cell-death was however significantly reduced compared to the *AvrPm17_THUN12* variant (paired Wilcoxon rank test p=0.0048) (Fig. 2D). We therefore concluded that *Pm17* can weakly recognise *AvrPm17_96224*, at least in a heterologous overexpression system. To address the question whether the weak recognition of *AvrPm17_96224* translates into phenotypes on *Pm17* wheat we made use of the above-mentioned transgenic lines Pm17#181 and Pm17#34 which were previously shown to exhibit differences in PM17 protein abundance with Pm17#181 representing the stronger line (16). Consistent with a prediction of a quantitative difference in AVR recognition, we observed reduced leaf coverage by mildew upon infection of the strong line Pm17#181 with isolate 96224 and with progeny of the 96224 X THUN-12 cross carrying the *AvrPm17_96224* haplotype (Fig. S1A-D), thus indicating that a residual recognition of *AvrPm17_96224* by transgenic *Pm17* overexpression is sufficient to reduce disease severity quantitatively. In contrast, the recognition of the *AvrPm17_THUN12* haplotype conferred complete disease resistance in both transgenic lines (Fig. S1A,B,D). Taken together these findings demonstrate that while the *Pm17* resistance in wheat genetically follows a classic gene-for-gene interaction model, phenotypic differences in *Pm17* mediated resistance is not only determined by sequence polymorphism in the AVR but also by Pm17 expression levels.

We therefore also analysed differences in AVR protein abundance using C-terminal FLAG epitope tagged AVRPM17 variants from Bgt_96224 and THUN-12. The presence of the FLAG epitope partially interfered with *AvrPm17* recognition since tagged versions exhibited significantly reduced HR levels when co-expressed with *Pm17*, however the specificity was not affected (Fig. S8A-D). Both AVRPM17-FLAG variants as well as PM17-HA were detectable on a Western blot (Fig. 2E, Fig. S8E-F). Interestingly, expression of AVRPM17_THUN12 in *N. benthamiana* resulted in higher protein abundance than for the weakly recognized AVRPM17-96224 variant (Fig. 2E). This suggests that the amino acid polymorphisms between 96224 and THUN-12 affect protein stability of AVRPM17 and therefore contribute to the differential recognition of the AVRPM17 variants by PM17, thus further demonstrating that protein expression levels of AVR and NLR variants are additional determinants underlying the seemingly binary gene-for-gene genetic determinism of immunity based on major R genes. Similar observations were recently described for *AvrPm3*^*a2/f2*^, where polymorphism in the AVR were found to affect protein amount and thereby directly influence recognition by *Pm3a* (32).

AVRPM17 is part of effector family E003, the second largest effector family found in *Blumeria graminis* (31). The family is comprised of small proteins of c.a. 110 amino acids that contain a predicted signal peptide, an N-terminal Y/FxC motif followed by a stretch of alternating hydrophobic residues as well as a conserved carboxy-terminal cysteine (Fig. S9). These features have been described for numerous *Blumeria* effectors, including all functionally characterized AVR proteins in wheat mildew (18, 20, 21). Using an *in silico* modelling approach based on IntFOLD5.0, we found that AVRPM17 is predicted to exhibit a ribonuclease-fold (pvalue = 1.145E-4), consisting of a single α-helix and three β-strands (Fig. S10A,B). A similar ribonuclease-fold was experimentally determined by crystallization for the barley powdery mildew (*B*.*g. hordei*) effector BEC1054 (33) and proposed for AVRA7, the avirulence gene of the barley NLR Mla7 (34). *Avra7* is part of the *AvrPm17* gene family (E003) (Fig. S11), suggesting that the ribonuclease-fold is conserved within the effector family. The particular arrangement of α-helix helix and β-strands has been predicted for other AVR proteins in wheat mildew. Most importantly such a pattern was described for the entire effector families E008, E018 and E034 encoding *AvrPm3*^*a2/f2*^, *AvrPm3*^*b2c2*^ and *AvrPm3*^*d3*^ respectively (18). We therefore hypothesize that *AvrPm17* and the *AvrPm3*’s encode for structurally similar proteins, despite little similarity on the primary amino acid sequences (Fig. S12) suggest that the wheat *Pm3* allelic series and its rye orthologue *Pm17* recognise structurally related effectors.

### *AvrPm17* is encoded in a mildew sub-lineage specific effector cluster and exhibits signs of re-occurring gene conversion events

Candidate effector genes in wheat and barley powdery mildew have been grouped into 235 families based on sequence similarity (31). *AvrPm17* belongs to family E003 which is represented with 69 members in *B*.*g. tritici* (isolate Bgt_96224), 70 members in *B*.*g. triticale* (isolate THUN-12) and 59 members in *B*.*g. hordei* (isolate *DH14*) ((31),(34)). E003 is physically organized in gene clusters distributed over seven of the eleven chromosomes of wheat and triticale mildew (Fig. S11). Family members encoded in the same chromosomal location form phylogenetically related clades, consistent with the previously proposed expansion mechanism of effector genes through local duplication ((22), Fig. S11). Interestingly, the *AvrPm17* clade is encoded in a gene cluster that spans more than 1.3Mb on chromosome 1 and contains 7 and 8 members in triticale and wheat mildew respectively, as well as a solitary family member on chromosome 8 (Fig. 3A,B). The locus also harbours an additional effector cluster from family E011, consisting of eleven members, that has expanded within the E003 cluster (Fig. 3B). To study the evolutionary history of the *AvrPm17* clade, we identified the region corresponding to the *AvrPm17* cluster on scaffold_27 in barley mildew isolate DH14 based on conserved syntenic flanking genes. Strikingly, the region in DH14 is only 200kb in size and harbours only two E003 family members of the *AvrPm17* clade as well as a single effector from family E011 (Fig. 3B). This indicates both effector clades have been significantly expanded in wheat mildew after the divergence from the barley mildew lineage (Fig. 3B). Using re-sequencing data from five rye mildew strains we could also show that rye mildew has six out of the eight E003 family members in the clade, whereas it lacks most of the E011 genes (Fig 3B), indicating that the E003 expansion has happened in a progenitor of rye/wheat mildew whereas the E011 family expansion in this region is wheat mildew specific. Strikingly, rye mildew does not encode for an *AvrPm17* gene, consistent with the virulent phenotype of the five rye mildew isolates on the *Pm17*-donor line ‘Insave’ (Fig. S13). This suggests that *AvrPm17* was lost in rye mildew likely due to the selection pressure imposed by the *Pm17* gene in the rye gene pool.

**Figure 3.**
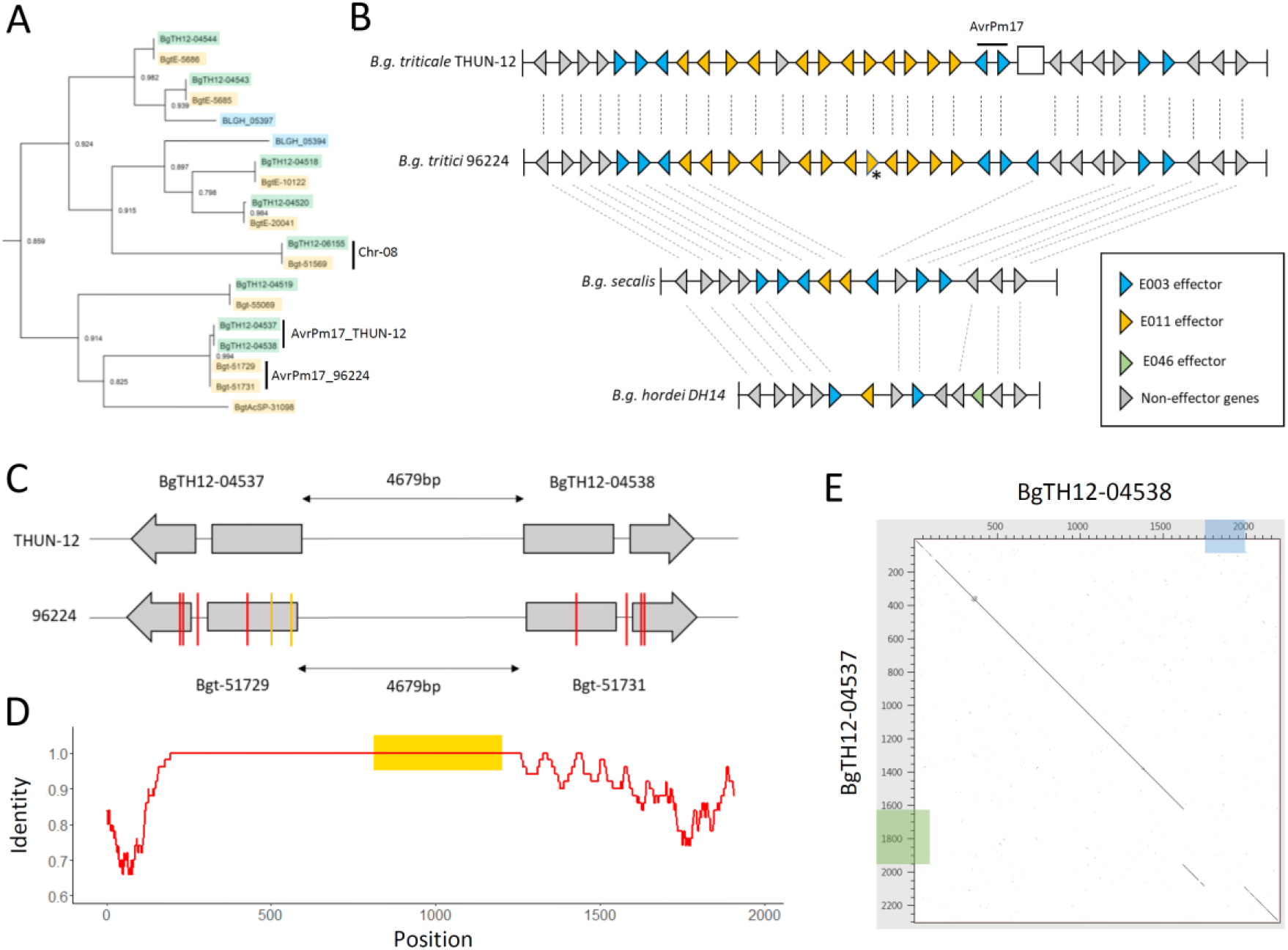
*AvrPm17* is a member of a highly expanded effector gene cluster. (A) Phylogenic relationship of the AvrPm17 effector family. Panel shows subsection of the phylogenetic tree based on protein sequences of E003 effector family members of *B*.*g. tritici* (69 members) *B*.*g*. triticale (70 members) and *B*.*g. hordei* (59 members). The full tree can be found in Fig. S12. Effector family members are highlighted as follows: members in *B*.*g. tritici* in yellow, members in *B*.*g. triticale* in green, and members in *B*.*g. hordei* in blue. For each branch, the local support values calculated with the Shimodaira-Hasegawa test are indicated. (B) Schematic representation of the *AvrPm17* effector cluster in the high-quality genomes of *B*.*g. triticale* THUN-12, *B*.*g. tritici* 96224, and *B*.*g. hordei* DH14. In the absence of a high-quality genome assembly for *B*.*g. secalis*, presence/absence of genes was estimated by coverage analysis based on mappings of five re-sequenced isolates. Genes that are present in at least one *B*.*g. secalis* isolate, where considered as present. Genes and their orientation are indicated by triangles. The white rectangle in the *B*.*g. triticale* THUN-12 assembly indicates the position of the 50kb deletion presented in Fig. 1F. The gene marked with an asterisk represents a collapsed gene duplication in the *B*.*g. tritici* 96224 assembly that was resolved in the *B*.*g. triticale* THUN-12 genome assembly. Syntenic relationship is indicated by dashed lines. The figure is not drawn to scale. (C-E). The AvrPm17 gene copies have evolved through gene conversion. (C) Analysis of SNPs in the *AvrPm17* gene copies in the two parental isolates. SNPs in THUN-12 are shown in comparison to 96224 for which both gene copies are identical. The *AvrPm17* genes are represented schematically with grey boxes representing the two exons. The transcriptional orientation is indicated by the direction of the arrowhead in the second exon. SNPs are indicated in the coding sequences and the intron, as well as in regions 100bp up- and downstream of the gene. Red bars represent SNPs that are shared between the two gene copies in THUN-12 and yellow lines indicate SNPs that are present in only one copy. (D) Visual representation of the duplication of *AvrPm17* in isolate 96224. To allow alignment of the two sequences, the insertions in the downstream region of the two genes were spliced out. The x-axis shows the alignment position, while the y-axis shows the sequence identity calculated in 50bp sliding windows. The position of the *AvrPm17* gene is highlighted by a yellow box. (E) Dotplot alignment of the duplicated gene copies and flanking region in the isolate THUN-12. Insertions in the downstream region of BgTH12-04537 and BgTH12-04538 are highlighted in green and blue, respectively.

Given the highly dynamic genomic context of the *AvrPm17* locus, it is striking that the avirulent AVRPM17_THUN12 variant and the partial gain-of-virulence variant AVPM17_96224 are encoded by near identical (*BgTH12-04537/BgTH12-04538*, two synonymous SNPs) or identical (*Bgt-51729/ Bgt-51731)* paralogous gene copies within the isolates THUN-12 and 96224, respectively (Fig. S14). Most importantly, the three non-synonymous SNPs that differentiate *AvrPm17_96224* from the avirulent *AvrPm17_THUN12* are identical in both genes *Bgt-51729 and Bgt-51731* (Fig. 3C). Congruently there is an identical SNP in the intron of *Bgt-51729 and Bgt-51731*. It is highly unlikely that the exact same four mutations have occurred independently in both genes, indicating a recent gene duplication in each isolate. However, the duplicated region in Bgt_96224 and THUN-12 is identical in size and position and therefore must have occurred in the ancestor of the two isolates (Fig. 3C). Consistent with the hypothesis of a more ancient duplication event we found that flanking regions of the duplication are significantly more divergent and contain two insertions (Fig. 3D/E, Fig. S15,16, Supplementary Text 1). These findings further demonstrate that the duplication is older than estimated based on the highly similar genic sequences. Therefore, the nucleotide polymorphisms defining the differences between *AvrPm17_THUN12* and *AvrPm17_96224* have most probably occurred in one gene copy and were then transferred to its duplicate by gene conversion (for further details see Supplementary Text 1). We propose that gene conversion event(s) contribute to the evolutionary potential of *AvrPm17* as an efficient way to transfer beneficial mutations to both gene copies.

### Virulent *AvrPm17* haplovariants are ancient and predate *Pm17* introgression into wheat

A haplotype mining approach in a diversity panel of 160 re-sequenced isolates of wheat mildew (138) and triticale mildew (22 isolates) for *AvrPm17* revealed there are three dominant AVRPM17 variants in the gene pool (Fig. 4A,B). Two of these are the above-described AVPM17_THUN12 (varA) and the weakly recognized AVPM17_96224 (varB). The most frequent haplotype is varC that contains a single amino acid polymorphism (A53V) and induces weaker HR in the *Nicotiana* co-expression assays as compared to the functional varA found in THUN-12 (Fig. 4C). In addition, eleven isolates originating from China encode for the only complete loss-of-recognition haplotype found (varD) with three amino acid changes (A53V, E55R, G61A) compared to varA (Fig. 4A-C, Fig. S17). Strikingly, 93% of the isolates with two *AvrPm17* copies encode for two identical mature AVRPM17 proteins in one of the following combinations; varA/varA, varB/varB and varC/varC (Fig. 4B). This supports the hypothesis that the *AvrPm17* gene copies are kept identical by recurring gene conversion events (for details see Supplementary Text 2, Fig. S14-16, S18-19, Table S2).

**Figure 4.**
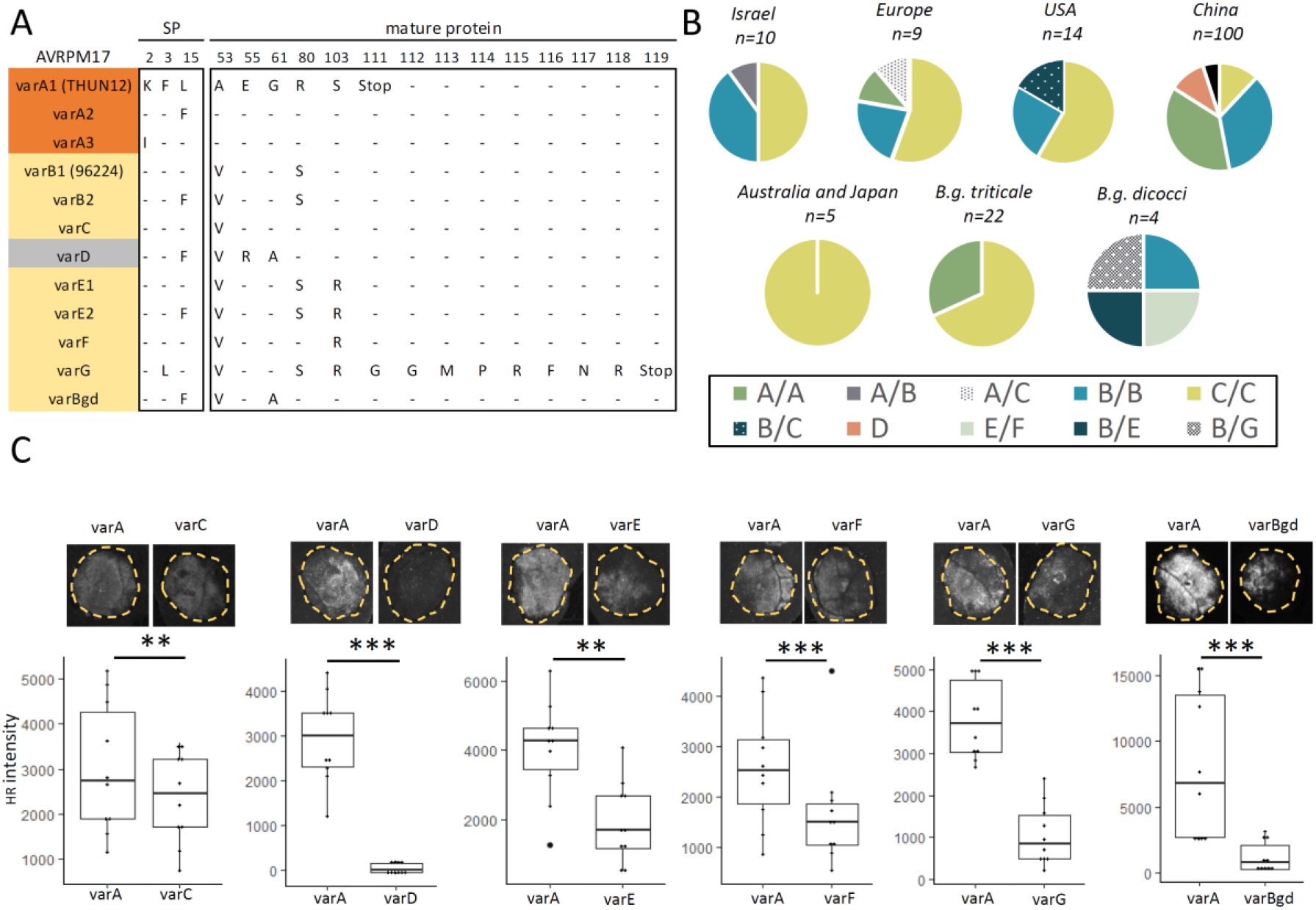
Diversity analysis of AVRPM17 in different *formae speciales*. (A) Protein variants found in a global mildew population of 160 isolates (*B*.*g. tritici, B*.*g. triticale, B*.*g. dicocci, B*.*g. dactylidis*). Colors indicate differences in recognition strength determined by transient co-expression in *N. benthamiana*. Orange indicates the avirulence allele varA and haplotypes that are recognized to a comparable extent. Yellow indicates variants that are recognized significantly weaker than the avirulent allele *AvrPm17*_varA and therefore represent partial gain-of-virulence alleles. The only full loss-of-recognition variant is indicated in gray. (B) Distribution of haplotypes in global *B*.*g. tritici, B*.*g. triticale* and *B*.*g. dicocci* populations. (C) Recognition strength of AVRPM17 variants (depicted in panel (A)) in *N. benthamiana* compared to AVRPM17_varA. Infiltrations were performed at R: Avr ratios of R: Avr of 1:1 and significance was assessed using a paired Wilcoxon rank sum test, significance level are indicated above the boxplots as follows: **=p<0.05, *** p<0.01.

We selected a set of 16 representative isolates covering the diversity of *AvrPm17* in wheat mildew (varA-varD), verified *AvrPm17* expression during early stages of infection (i.e. haustorial stage, Fig. S20) and analyzed their virulence phenotype on *Pm17* transgenics. Consistent with the recognition strength in *N. benthamiana*, representative isolates encoding for *AvrPm17_varD* were fully virulent and isolates carrying *varA*, with the exception of one isolate, were avirulent on the transgenic *Pm17* lines (Table S3). Isolates carrying the weakly recognized variants *AvrPm17_varB* and *AvrPm17_varC* displayed intermediate phenotypes on the transgenic lines (Table S3). Thus, the recognition strength of *AvrPm17* haplovariants observed in *N. benthamiana* largely correlated with disease resistance in wheat thereby confirming the biological relevance of the heterologous *Nicotiana* system to study quantitative effects of *Avr* recognition by resistance genes. Furthermore, these findings indicate that *AvrPm17*_varD and *AvrPm17*_varB/varC indeed represent virulence or partial virulence alleles respectively, that are likely responsible for the resistance breakdown of the *Pm17* gene in wheat.

Both partially virulent variants varB and varC were present in all major subpopulations (i.e. China, Europe, Israel, USA) (Fig. 4B). Given their global distribution and considering that some of the isolates were collected already in the 1990’s (SI appendix Dataset 3) this suggests that the partially virulent *AvrPm17* variants varB and varC were present as standing genetic variation in the wheat mildew population before large-scale agricultural deployment of wheat varieties carrying the *Pm17* introgression at the beginning of the 21^st^ century in the US, and only subsequently in other regions of the world (27, 29). To further test this hypothesis, we extended our haplotype analysis to closely related *formae speciales* of *B*.*g. tritici*. Due to their distinct host range, *B*.*g. dicocci*, a *forma specialis* sampled on wild tetraploid wheat and *B*.*g. dactylidis* infecting the wild grass *Dactylidis glomerata* (23, 35) are unlikely to have previously been exposed to the *Pm17* resistance gene. Strikingly, we found that three isolates of *B*.*g. dicocci*, encode up to two copies of *AvrPm17_varB* (Fig. 4B). Furthermore, we found three additional haplovariants (varE-G) specific to *B*.*g. diccoci*. Co- expression of varE-G with *Pm17* in *N. benthamiana* resulted in significantly weaker HR responses compared to *AvrPm17_varA*, indicating these variants also represent partially virulent alleles (Fig. 4C). This is consistent with the observation that these haplovariants share the A53V, R80S mutation (varE,G) or the A53V mutation (varF) with AVRPM17_varB (Fig. 4A). Since *B*.*g. dicocci* does not grow on most hexaploid wheat cultivars, including ‘Bobwhite’ (23) we could not test the contribution of the varE-G recognition to *Pm17* virulence. In *B*.*g. dactylidis*, represented by two isolates, we found an additional haplovariant *AvrPm17*_varBgd which carries two substitutions (A53V and G61A) compared to *AvrPm17_varA* and is only very weakly recognized by *Pm17* in *N. benthamiana* (Fig. 4A/C). Most importantly, these mutations are shared with the non-recognised Chinese haplotype *AvrPm17_varD*, demonstrating that (i) the E55R substitution in the Chinese haplotype is the causative mutation leading to complete loss-of-recognition by *Pm17* (Fig. 4C, Fig. S17) and (ii) that part of the *AvrPm17* diversity found in wheat mildew is ancient and predates the split of *B*.*g. tritici* and *B*.*g. dactylidis*. Taken together we found that a significant proportion of the *AvrPm17* sequence diversity found in *B*.*g. tritici*, including several gain of virulence mutations, is shared with its closely related *formae speciales B*.*g. dicocci* or *B*.*g. dactylidis*. Combined with the observation of a global distribution of partially virulent *AvrPm17* variants B and C and their presence in isolates collected before the deployment of *Pm17* wheat in agriculture, our findings strongly indicate that these *AvrPm17* gain of virulence mutations represent standing genetic variation in wheat mildew which predates precedes the introgression of *Pm17* into wheat.

While the existence of numerous virulent or partially virulent *AvrPm17* haplotypes in the global mildew population prior to *Pm17* introgression might explain rapid *Pm17* resistance breakdown this observation is hardly compatible with the initially described broad resistance phenotype exerted by the 1AL.1RS translocation. Based on these considerations we therefore propose the 1AL.1RS translocation to harbor a second mildew resistance gene in addition to *Pm17*.

### The 1RS.1AL translocation encodes for two powdery mildew resistance specificities

To test for the predicted second resistance gene of the 1AL.1RS translocation, we characterized the genetic association of avirulence of the 96224 X THUN-12 mapping population on the original 1AL.1RS translocation line ‘Amigo’ (16). The *Pm17* avirulent isolate THUN-12 showed an intermediate phenotype on ‘Amigo’, demonstrating that recognition of *AvrPm17_varA* results in quantitative resistance in presence of the endogenous *Pm17* gene (Fig. 5A,B). In contrast, the *Pm17*-virulent isolate 96224 was avirulent on ‘Amigo’, indicating that (i) this isolate carries an additional avirulence component recognized by ‘Amigo’ and (ii) the 96224 X THUN-12 bi-parental population is suited to validate the second resistance specificity in ‘Amigo’ (Fig. 5C). Consistent with our hypothesis, a QTL mapping analysis based on 117 progeny of Bgt_96224 X THUN12 identified two significant QTLs associated with the avirulence phenotype on cultivar ‘Amigo’ (for details see Supplementary Text 3, Fig 5D, Table S4). One QTL on chromosome 1 corresponds to the *AvrPm17* locus thereby verifying the activity of the *Pm17* gene in the original translocation line ‘Amigo’. In addition, we identified a highly significant QTL on chromosome 9 that was not detected in the QTL analysis on the transgenic *Pm17* lines (Fig. 1A, Fig. 5D), likely encoding the avirulence component recognized by the predicted second resistance gene of the 1AL.1RS translocation. The confidence interval of the QTL on chromosome 9 encompasses 371 kb in the avirulent isolate 96224 and harbors a total of 16 effector genes of which four are polymorphic compared to THUN- 12 (Table S5, Fig. S21A). Upon co-expression with *Pm17* in *N. benthamiana* none of the effector candidates encoded by isolate Bgt_96224 within the confidence interval elicited a hypersensitive response (Fig. S21B). This finding demonstrate that the second QTL on chromosome 9 is independent of the *Pm17* resistance specificity. Most importantly, only progeny of the cross that carry the *AvrPm17_96224* haplovariant (*AvrPm17_varB*) and the THUN-12 genotype in the QTL on chromosome 9 are fully virulent (Fig. 5D,5E), further demonstrating that the simultaneous presence of both virulence alleles is necessary to overcome the resistance on ‘Amigo’.

**Figure 5.**
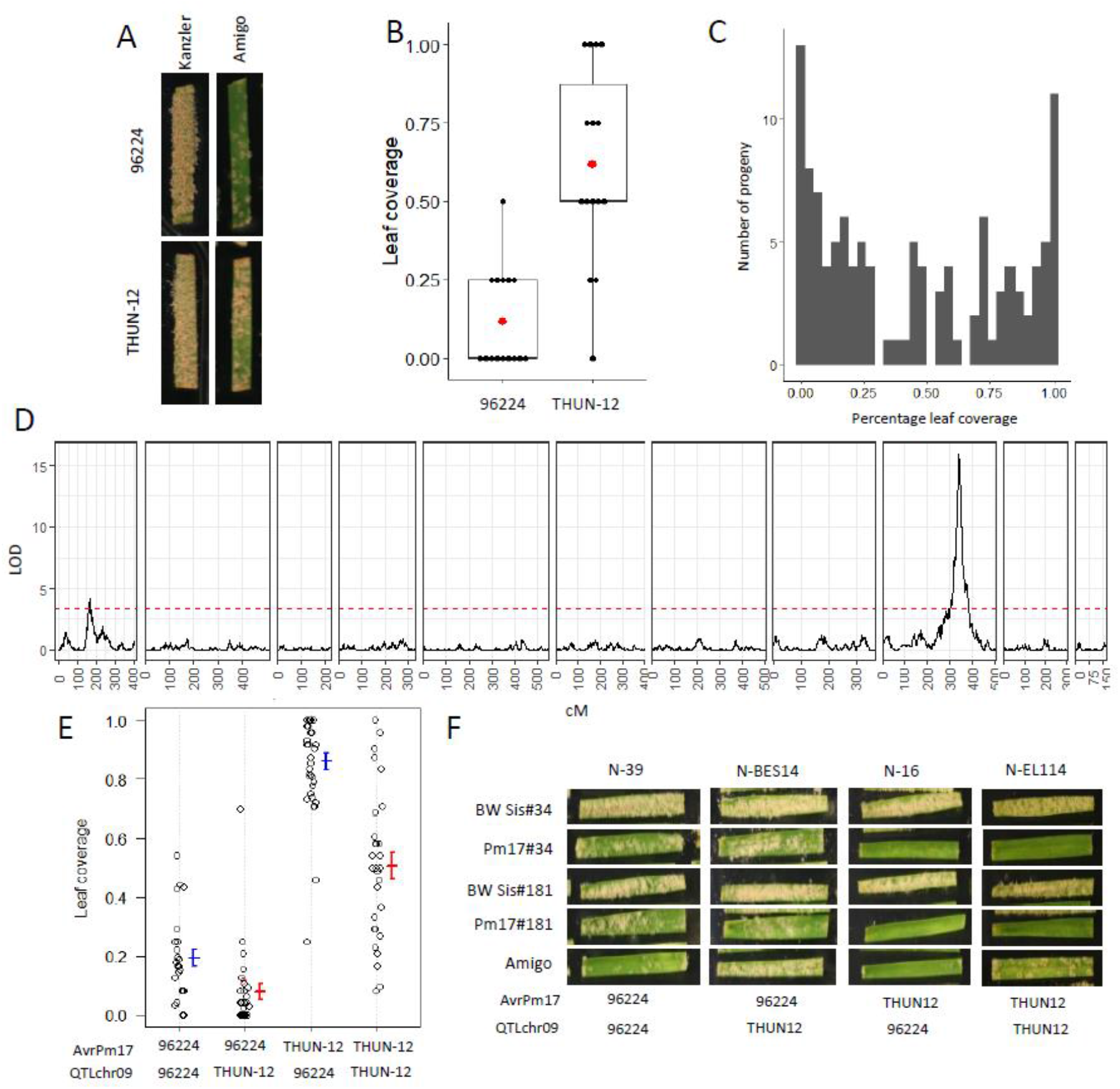
QTL mapping in the Bgt_96224 X THUN-12 F1 population on the wheat cultivar ‘Amigo’ carrying a 1RS.1AL translocation from ‘Insave’ rye including the *Pm17* resistance gene. (A) Representative photographs of phenotypes of the parental isolates Bgt_96224 and THUN-12 on Amigo at 10dpi. The susceptible wheat cultivar ‘Kanzler’ was used as an infection control. (B) Boxplot summarizing the phenotypes of Bgt_96224 and THUN-12 on ‘Amigo’. Leaf coverage of individual leaf segments was scored according to the following scale: avirulent = 0, avirulent/intermediate 0.25, intermediate 0.5, intermediate/virulent =0.75, virulent = 0. (C) Distribution of phenotypes of the 117 progeny of the cross Bgt_96224 X THUN-12 on ‘Amigo’. Progeny phenotypes were scored as described in (B) and the average of at least 6 leaf segments for each progeny was plotted (D) Single interval QTL mapping of Bgt_96224 X THUN-12 on cultivar ‘Amigo’. The black line indicates the LOD (logarithm of the odds) score of the association throughout the 11 chromosomes of wheat powdery mildew. Red line indicates the significance threshold determined by 1000 permutations. (E) QTL effect plots summarizing phenotypes of the 117 progeny on ‘Amigo’. The phenotypes were plotted based on the genotypes of the best associated marker at the QTL location on chromosome 9 (QTLchr09) and chromosome 1 (AvrPm17 locus). (E) Photographs of representative progeny of the cross Bgt_96224 X THUN-12 with different genotype combinations (see E) for the QTLs identified in (D). Phenotypes on wheat cultivar ‘Amigo’ and the transgenic lines expressing *Pm17* at 10dpi are shown.

We therefore conclude that the broad effectiveness of the 1AL.1RS translocation in providing resistance against wheat powdery mildew was based on two resistance gene specificities in the 1AL.1RS translocation. Since the *Pm17* resistance specificity has been attributed to a single locus based on segregation analysis we hypothesize that the second locus is genetically linked on the 1RS.1AL region and has previously been genetically masked due to repressed recombination frequently associated with introgressed segments in wheat (36).

In summary, we here demonstrate that the *Pm17* introgression is genetically complex and that such complexity could only be revealed through accurate genetic dissection of the avirulence determinants in the pathogen distinguishing the two resistance specificities.

## Discussion

The recent identification of numerous *Avr* genes both in *B*.*g. tritici* and *B*.*g. hordei* has significantly advanced our understanding of NLR mediated resistance in the cereal powdery mildew pathosystem (17, 18, 20-22, 34, 37). The functional cloning of *Avr* genes not only allowed molecular studies on recognition mechanisms (32, 34) but has also set the ground for genetic studies based on the natural diversity of avirulence components in local and global mildew collections. This has led to the discovery of numerous gain of virulence mechanisms exerted by *Blumeria* pathogens, including single amino acid polymorphisms, truncations and deletions of *Avr* genes as well as a fungal encoded suppressor *SvrPm3* acting on *Pm3* mediated resistance through masking of AVRPM3 recognition (17, 18, 34). These findings highlight the importance of genetic and genomic studies in fungal plant pathogens in order to understand the mechanisms of resistance breakdown and allow us to adapt current breeding approaches towards more durable deployment of resistance genes in cereal crops.

With so far ten *Blumeria Avr* genes cloned and functionally characterized several patterns emerged. *Blumeria* AVR effectors were found to be small proteins with a length of 102-130 amino acids, to contain an N-terminal signal peptide, a largely conserved Y/FxC motif and a conserved cysteine residue towards the C-terminus (17, 18, 20-22, 34, 37) while otherwise exhibiting highly divergent amino acid sequences. Furthermore, wheat mildew *Avrs* were consistently among the highest expressed genes within their effector gene family, indicating high abundance in host cells upon secretion presumably influencing the efficacy of their virulence function alongside NLR mediated recognition in resistant cultivars (18). The newly identified *AvrPm17* exhibits all the above-mentioned characteristics of *Blumeria* AVRs and therefore further corroborates the emerging patterns.

Despite showing little homology to proteins with a known function, more than a hundred *Blumeria* effectors are predicted to exhibit a ribonuclease-like fold (20, 33, 34, 38, 39). Notably, such a ribonuclease-like structure has recently been confirmed by protein crystallization of the barley powdery mildew effector BEC1054 (33). Similarly, despite highly divergent amino acid sequences, *in silico* protein modeling approaches predicted ribonuclease-like folds for most of the functionally verified avirulence proteins in barley and wheat mildew (20, 21, 34), including all AVRPM3 effectors (18) and AVRPM17 (this study). Based on the homology between rye *Pm17* and wheat *Pm3* NLRs combined with the predicted structural similarities of their corresponding AVR proteins we propose a conserved recognition mechanism, likely leading to similar selection pressures acting on AVR genes for evasion of recognition.

Extensive haplovariant mining in a global wheat mildew collection for *AvrPm3*^*a2/f2*^, *AvrPm3*^*b2/c2*^ and *AvrPm3*^*d3*^ revealed that virulent alleles were exclusively based on single amino acid polymorphisms (18, 22). Even though copy number variation was common, disruption or deletion of the avirulence gene, as observed for many other *AVR*s has never been detected. In line with these findings, our haplovariant mining approach for the paralogous *AvrPm17* copies in a comparable wheat mildew diversity panel also failed to identify gene deletions or non-sense mutations and in return found four variants varA-varD of which three represent partial or complete virulence alleles that are based on amino acid polymorphisms. In contrast, we found the *AvrPm17* genes deleted in rye mildew, suggesting different gain of virulence mechanisms in these closely related mildew sub-lineages, likely due to differences in the exposure to the *Pm17* gene.

Gene duplications in effector genes are common and considered advantageous for pathogens as they allow the independent diversification of virulence factors (31). However, the presence of identical avirulence gene copies can represent a major liability as gain of virulence mutations need to occur in both gene copies to effectively change the phenotypic outcome. This was described for wheat mildew *AvrPm3*^*d3*^ in which gain of virulence mutations in one of the tandem duplicated gene copies was not sufficient to render the isolates virulent (18). Gene conversion, efficiently transferring beneficial mutations between gene copies, could provide pathogens with a molecular mechanism to mitigate the disadvantage of duplicated avirulence genes. Indeed, a case of gene conversion leading to gain of virulence was described for *Avr3c* in the oomycete *Phytophthora sojae*, (40). Similarly, we found evidence for gene conversion to have occurred between the paralogous copies of *AvrPm17* (Supplementary Text 1 & 2). The high frequency of wheat mildew isolates encoding for varA/varA, varB/varB or varC/varC genotypes furthermore indicates repeated gene conversion events between the two paralogs. Whether this phenomenon is dependent on inherent predisposition of the locus to non-allelic gene conversion to occur or whether it reflects the existence of an additional selection pressure linked to the virulence function of *AvrPm17* to maintain the sequences identical will be subject to further studies.

It was hypothesized that introgressed *R* resistance genes provide effective and durable resistance by recognizing an effector gene which, in the absence of previously acting diversifying selection, is largely conserved (7). The opportunity to test this hypothesis for a fungal pathogen in wheat arose with the recent cloning of the introgressed stem rust resistance genes *Sr35* (from *T. monococcum*, (41)) and *Sr50* (from *Secale cereale*, (42)) and their corresponding avirulence genes *AvrSr35* and *AvrSr50* in *Puccinia graminis* f. sp. *tritici (Pgt*) (43, 44). Virulent alleles for both genes were identified in *Pgt* races with diverse geographic origins. Whether these virulent alleles emerged before the introgression of *Sr35* and *Sr50* into wheat or as a consequence of their agricultural use was however not assessed. The identification of partially or fully virulent *AvrPm17* haplovariants (varB- varD) in a geographically diverse set of wheat mildew isolates collected over the last three decades and the identification of *AvrPm17* homologs in closely related mildew sublineages provided a unique opportunity to investigate a possible connection between the starting agricultural use of the *Pm17* introgression in wheat at the beginning of the 21^st^ century and the emergence of virulent *AvrPm17* alleles in wheat mildew. Using (i) a global mildew population with a unique temporal resolution including many isolates that were collected before *Pm17* deployment or exhibit different host preferences (ii) careful phenotypic studies on transgenic *Pm17* lines, and (iii) functional studies of *Avr* recognition in transient protein expression assays, we could demonstrate that virulent *AvrPm17* variants were largely present in mildew populations prior to the deployment of *Pm17*. We propose that this genetic diversity has arisen from the evolutionary arms-race between *Blumeria* and its host species potentially tracing back to a Pm17/Pm3-like gene in the progenitor of rye and wheat. This hypothesis is corroborated by the fact that the *AvrPm17* gene is encoded in a highly expanded gene cluster of effector family E003, which is exclusive to wheat and rye mildew, suggesting that the expansion of this cluster evolved prior to the split of the two mildew lineages 250’000 years ago (23). One of the mechanisms that is proposed to drive expansion of effector gene clusters is the continuous coevolution with the host immune system (45, 46). Thus, the presence of *Pm17/Pm3*-like genes in the progenitor of rye and wheat might have resulted in selection pressure leading to the expansion of the effector cluster on chromosome 1 in the progenitor of wheat and rye mildew, suggesting a long history of *R-*gene mediated effector evolution in natural ecosystems, long before the start of agricultural cultivation. Our findings highlight the importance for resistance durability to select introgressed resistance specificities based on the evolutionary history of donor and recipient species. In this work, we demonstrate the necessity to identify and monitor the genetic diversity of the corresponding avirulence factors in order to achieve effective and durable resistance. We propose that such studies are very timely, considering the current important efforts to introgress *R* genes into wheat from phylogenetically distant wild relatives or phylogenetically close diploid progenitor species.

An often-stated advantage of larger translocations from related species is the simultaneous introgression of several resistance specificities active against different plant pathogens such as the most widely deployed rye translocation 1BL.1RS from ‘Petkus’ carrying *Lr26, Yr9, Sr31* and *Pm8* (4). For effective and durable resistance, the introgression of several resistance genes active against the same pathogen is highly desirable. By extending the mildew QTL mapping approach from *Pm17* transgenic lines to the original 1RS.1AL translocation cultivar ‘Amigo’ we have found evidence for the presence of a second resistance gene potentially recognizing an avirulence gene of *B*.*g. tritici* isolate Bgt_96224. Historically the *Pm17* resistance associated with the 1RS.1AL translocation has been attributed to a single locus (47). The additional resistance specificity predicted by our QTL approach is therefore most likely genetically linked with the *Pm17* gene and has been missed by genetic approaches solely applied on the plant side due to suppressed recombination within the translocated genomic region originating from ‘Insave’ rye (36). The simultaneous presence of two race-specific resistance genes in the 1RS.1AL translocation might explain the initially broad resistance exhibited by cultivars such as ‘Amigo’, despite the likely long-standing presence of several gain of virulence alleles for *AvrPm17* in the *B*.*g. tritici* population. The identification of this second so far unknown AVR/R gene pair in the future will potentially provide further answers regarding the initial efficacy but also the quick breakdown of the powdery mildew resistance encoded on the 1RS.1AL translocation.

Identification and cloning of introgressed resistance genes has often been hampered by the absence of recombination throughout parts or the entirety of the alien chromatin regions (48). In recent years several technological advances such as RenSeq or MutChromSeq approaches that do not rely on fine-mapping have helped to alleviate this phenomenon and led to the identification of numerous new resistance genes often residing in highly complex loci (49, 50). Here we show that genetic mapping populations of plant pathogens could provide an additional tool to dissect complex translocated genomic regions with low or absent recombination thereby complementing recently developed, plant-focused approaches.

## Materials and Methods

Detailed Material and Methods section is available as SI appendix. Constructs used in this study are listed in SI appendix Dataset 1. Primer sequences are listed in SI appendix Dataset 2. Details about powdery mildew isolates and their associated (SRA) accession numbers are listed in SI appendix Dataset 3. Phenotyping and subsequent QTL analysis are described in SI appendix section 1. Candidate identification is described in SI appendix section 2 Construction of the expression plasmids are described in SI appendix section 3. Transient expression procedure using *Agrobacterium tumefaciens* in *Nicotiana benthamiana* followed by hypersensitive response measurement are described in SI appendix section 4, western blot detection of tagged avirulence and resistance genes can be found in SI appendix section 5. Expression analysis can be found in Si appendix section 6. Bioinformatic analysis are detailed under SI appendix section 7. The Sequence of *Pm17* is available at GeneBank under the accession number AYD60116.1. AvrPm17 haplovariants are available under the accession numbers XX-XX.

## Acknowledgments

This work was supported by the University research priority program (URPP) “Evolution in Action” of the University of Zürich, Switzerland and the Swiss National Science Foundation grant number 310030B_182833. The authors would like to thank Prof. Daniel Croll from the University of Neuchâtel for helpful input on the topic of gene conversion.

