## Supplementary_Material for "Standing genetic variation of the AvrPm17 avirulence gene in powdery mildew limits the effectiveness of an introgressed rye resistance gene in wheat"

**This PDF file includes:**

Supplementary Text 1-3
Material and Methods
Figures S1 to S21
Tables S1 to S5
Legends for Datasets S1 to S3

**Other supplementary materials for this manuscript include the following:**

Datasets S1 to S3

**Supplementary Text**

**Supplementary Text 1: Recurrent events of gene conversion underline the evolution of the *AvrPm17* locus**

Gene duplication followed by sequence diversification has been suggested as a driving force to increase genetic diversity (1). The *Blumeria graminis* effector complement is characterized by high rates of gene duplication (2). In line with this finding, both isolates Bgt_96224 and THUN-12 contain a duplicated region of 2'300 bp of which 386bp encode *AvrPm17*. The high identity of the two copies of the coding sequences within the isolates (100% in Bgt_96224, 99% in THUN-12) indicated that the duplication is recent. However, when the analysis was extended to the entire duplicated region (813bp upstream and 1100bp downstream both genes) we found that the sequences have considerably diverged (Fig 3D, Table S2, Fig. S15). The distal 200bp segment of the 2'300 bp long duplication shares 79% and 84% identity upstream and downstream respectively (Table S2). In addition, each of the two duplicated regions carries a specific insertion that is not present in the other duplicate. (Fig. 3E, S16). This high divergence in the flanking regions indicates that the duplication is older than estimated based on gene identity. A mechanisms that best explains this structure is gene-conversion, a phenomenon that leads to non-reciprocal exchange of DNA between homologous sequences (3). Experimentally determined size of gene conversion (gene conversion tracts) in yeast are generally small (2kb) (4), however due to the small size of effector genes (<500bp), entire genes can be homogenized by gene conversion. In the case of AvrPm17, the data suggest the locus has undergone re-occurring gene conversion events between the two paralogous copies probably due to selection pressure to maintain the middle part of the duplication, containing the open reading frame, identical (Fig. 3D).

The genic SNP-pattern in the *AvrPm17* genes revealed by the two high quality genome sequences of *B.g. triticale* THUN-12 and *B.g. tritici* 96224, provides further evidence for gene-conversion events. Considering the avirulent THUN-12 as reference, both gene copies in Bgt_96224 have the same four SNPs (three in exons and one in the intron) (Fig. 3C, S14). The duplications in both Bgt_96224 and THUN-12 are however identical in size and position, indicating that the duplication must have existed in the ancestor of the two isolates. The most likely scenario is that the SNPs occurred in one gene copy and were then transferred to the second copy trough gene conversion. This possibly represents a relatively recent non-allelic gene-conversion event as the two gene copies as well as their 100 bp up- and 100 bp downstream region show complete sequence identity within the otherwise more diverged duplicated region. Thus, we propose that non-allelic gene conversion played an important role in the *AvrPm17* evolution, by transferring beneficial mutations that occurred in one copy to its duplicate. Such a case has previously been described in *Phytophthora sojae*, where loss-of-recognition occurred in one copy of the tandemly duplicated *Avr3c* genes through amino acid substitution and was subsequently spread to the second gene copy via gene conversion (5). Gene duplications are considered advantageous for pathogens as they allow the independent accumulation of mutations and subsequent diversification of virulence factors (6). However, in case of recognition by a corresponding immune receptor the presence of identical avirulence gene copies represents a major disadvantage as gain of virulence mutations need to occur in both gene copies to escape recognition. This is exemplified by the tandemly duplicated *AvrPm3*$^{d3}$ gene pair, in which one gene copy encodes amino acid polymorphisms leading to loss-of-recognition by *Pm3d* in numerous *B.g. tritici* isolates (7). However, the strains remain avirulent on *Pm3d* wheat lines due to the presence of a second avirulent *AvrPm3*$^{d3}$ version. Thus, the spread of gain-of-virulence mutations from one gene copy to the second copy could represent a mechanism to cope with the adverse effect of encoding for multiple paralogous avirulence gene copies.

**Supplementary Text 2: The ancestor of *B.g. tritici* encoded two *AvrPm17* copies**

Using genomic mapping coverage as a proxy (2), we estimated that 90% of the global 164 *B.g. tritici* and *B.g. triticale* isolates contain two *AvrPm17* genes. The only exceptions were the 11 isolates from China encoding *AvrPm17_varD* and an additional three isolates that likely encode more than two copies of *AvrPm17* (Fig. S19). Strikingly, 93% of the isolates with two *AvrPm17* copies encode for identical mature proteins in one of the following combinations; varA/varA, varB/varB and varC/varC (Fig. 4B). These findings indicate recurring gene-conversion events between *AvrPm17* paralogs and indicate the presence of a selection pressure, possibly linked to effector function in virulence, to maintain the sequences identical.

To determine if the isolates with one *AvrPm17* gene copy represent the ancestral state in wheat mildew, we compared the gene copies in Bgt_96224 and THUN-12 with isolate GZ-6 carrying the *varD* haplotype and *B.g dactylidis* that both contain a single *AvrPm17* gene. The 100bp sequence upstream of the start codon of *BgTH12-04537* contains nine SNPs compared to the corresponding region of *BgTH12-04538* and *Bgt-51729/Bgt-51731*; whereas the 100bp downstream are identical in all four genes (Fig S.14). Interestingly, eight of the nine SNPs in the polymorphic 100bp upstream region of *BgTH12-04537* are found in the same region of *AvrPm17_Bgd* in *B.g. dactylidis* and of *AvrPm17_varD* in isolate GZ-6 (Fig S.14). Minimal parsimony assumptions would suggest that isolates with a single *AvrPm17* gene represent the ancestral state prior to the duplication. However, alignment of the duplicated regions in Bgt_96224 and THUN-12 with the corresponding region in isolate GZ-6 (encoding varD) shows that the first 200 bp of the region in GZ-6 is identical to the first gene copy (*BgTH12-04537/Bgt-51729)* (100% and 99.5% identity, respectively, Table S2) whereas the 200 bp end of the segment is more similar to the second gene copy (*BgTH12-04538/Bgt-51731*) (90% identity compared to 86%) (Figure S15, Table S2). In addition, the downstream region of the gene in GZ-6 contains the same insertion as *BgTH12-04538/Bgt-51731* and lacks the insertion in *BgTH12-04537/Bgt-51729* (Fig. S18). This suggests that isolates containing a single *AvrPm17* gene represent a derived state that lost one copy possibly through recombination of the two paralogs. In summary, our data strongly suggest that the ancestor of *B.g. tritici* already encoded two *AvrPm17* paralogs that exchanged mutations through gene conversion.

**Supplementary Text 3: QTL mapping of the *B.g. tritici* biparental F1 population derived from a cross of Bgt_96224 X THUN-12 on wheat cultivar 'Amigo'**

We predicted the presence of an additional *R* gene in the original 1RS.1AL translocation line 'Amigo' which partially or completely masks the effect of *Pm17* (8). We therefore phenotyped 117 progeny of Bgt_96224 X THUN-12 on cultivar 'Amigo' and observed highly quantitative segregation of the progeny (Fig. 5C). Subsequent single interval QTL mapping identified two significant QTLs, together explaining 75% of the phenotypic variance observed on cultivar 'Amigo' (Fig. 5D, Table S4). The first QTL is located on chromosome 9 at 340cM (LOD=15.97) whereas the second QTL locates to chromosome 1 at 165.8cM (LOD=4.24) which corresponds to the *AvrPm17* locus found on transgenic *Pm17* lines (Fig. 5D, Fig. 1A) thereby confirming the activity of *Pm17* in 'Amigo'. Since the highly significant QTL located on chromosome 9 was previously not found on *Pm17* transgenics, we hypothesized that it harbors an additional avirulence gene recognized by the predicted second resistance specificity active against *B.g. tritici* in the 1RS.1AL translocation of 'Amigo'. In this scenario the 'Amigo' avirulent isolate Bgt_96224 encodes for the avirulence component of this unknown resistance gene whereas THUN-12 encodes for the corresponding virulent allele. Indeed, segregation analysis of the progeny of Bgt_96224 X THUN-12 revealed that full virulence on 'Amigo' was specifically observed for progeny containing the *AvrPm17_96224* haplovariant and the THUN-12 genotype in the QTL on chromosome 9 (Fig. 5E,F).

The genetic confidence interval of the QTL on chromosome 9 (1.5LOD) delimited by markers snp106219 and snp106395 encompasses a physical interval of 371'034 bp in the *B.g. tritici* 96224 assembly (Table S5, Fig. S21A). The interval encodes 38 genes, of which 16 represent candidate effector genes encoding for 13 unique proteins (Fig. S21A, Table S5). Furthermore, two non-effector genes, *Bgt-3045* and *Bgt-3046*, contain a predicted signal peptide and are therefore likely secreted. The majority of the encoded candidate effectors in this locus (14 of 16) are part of candidate effector family E001, the largest candidate effector family in *Blumeria* (2). Interestingly, only five effectors are polymorphic between the isolates Bgt_96224 and THUN-12: *Bgt-51585*, Bgt-70025 and *BgtE-20010* contain one amino acid polymorphism in the mature peptide and *BgtA-21577* contains four. In addition, Bgt-70077 is duplicated in THUN-12 and the two proteins contain one and seven amino-acid polymorphisms compared to the 96224 protein, respectively (Table S5). To rule out the possibility that the QTL on chromosome 9 represents a second *Pm17* avirulence locus we co-expressed all 16 candidate effector genes as well as the presumably secreted *Bgt-3045* and *Bgt-3046* with *Pm17* in *N. benthamiana* (Table S5, Fig. 21B). None of the candidate genes triggered a hypersensitive response (Fig. S21B). Thus, we conclude that the QTL on chromosome 9 does not represent an avirulence locus corresponding to the rye resistance gene

*Pm17* but likely encodes an avirulence component recognized by a second *R* gene in the cultivar 'Amigo'.

In summary, we hypothesize that the original 1AL.1RS translocation present in cultivar 'Amigo' carries two *R* genes inherited from 'Insave' rye. This stands in contrast to previous reports which indicated a single powdery mildew resistance specificity (*Pm17*) in Amigo (9). However, the 1AL.1RS translocation has been described to display repressed rates of recombination (10) and to our knowledge, an analysis of individual mutants which would allow to dissect resistance specificities encoded by multiple linked *R* genes has never been performed on the 1AL.1RS translocation. Here we used a pathogen-based strategy using a segregating mildew population to decipher the two resistance specificities in 'Amigo'.

**SI appendix Material and methods**

**Section 1 Plant material, mildew isolates, phenotyping and QTL analysis**

Crossing, genotyping and genetic map production of the mapping population Bgt_96224 X THUN-12 was described in (2). F1 progeny were phenotyped on the *Pm17*-donor line 'Amigo' and *Pm17* transgenic lines BW Pm17#34 and BW Pm17#181 described in (8) and scored at 10 days post infection as described in (7). The susceptible cultivar 'Kanzler' was used as infection control. Phenotypes were assessed for individual leaf segments according to the following scale: avirulent = 0, avirulent/intermediate 0.25, intermediate 0.5, intermediate/virulent =0.75, virulent = 1 and final score consist of an average of at least four leaf segments.

Single interval QTL mapping was performed using the R/qtl v.1.46.2 (https://rqtl.org/) package in Rstudio (v1.2.1335, (11). The genetic map was processed using the commands read.cross(), jittermap() and calc.genoprobe(, step=1). Due to the non-normalized distribution of the phenotypes, single interval analysis was performed using the scanone(model="np") method. Significance levels were established with 1000 permutations of the scanone command. Genetic confidence intervals were extracted using the command lodint(expandtomarkers=TRUE). Percentage of the variance explained by the two QTL was calculated using the command sim.geno(step=1, n.draws=100, err=0.01) followed by the command makeqtl(, chr=c(9,1), pos=c(340,165.7)) and fitqtl(method="hk",pheno.col=2, formula=y~Q1+Q2+Q1:Q2).

**Section 2 Candidate identification**

*AvrPm17* candidate identification was based on the assembly and annotation of parental isolates Bgt_96224 (2) and THUN-12 (unpublished data). The physical intervals underlying the genetic confidence intervals were analysed manually, and correctness of the annotation was assessed using RNA-Seq data from both parental isolates (12, 13). For this purpose, RNAseq was mapped against the reference genome with STAR (v2.6.0a, (14)) using the method described in (15) and visualized with the integrative genome viewer IGV (v2.8.6, (16). Low quality gene models, overlapping with transposable elements, lacking transcriptional support or disrupted open-reading frames were excluded from the analysis. Erroneous gene models were corrected manually. Signal peptide prediction was performed with SignalP5.0 (17). Differential gene expression analysis was performed using EdgeR (v3.11, (18)) as described in (15). LogFold change (logFC) values of >1.5 was considered as significant.

**Section 3 Expression constructs**

The molecular identification of the *Pm17* gene and C-terminal epitope tagging with a hemagglutinin (HA) epitope have been described in (8). *Pm17*-HA was transferred into a gateway compatible entry plasmid using the pENTR-D-TOPO Kit (Invitrogen) according to the manufacturer. Transfer of *Pm17*-HA into the binary expression vector pIPKb004 (19) was achieved using Gateway LR clonase II (Invitrogen). Expression constructs of epitope-tagged *Pm3* alleles and *Pm8* have been previously described in (7).

A complete list of all effector constructs produced by gene synthesis by our commercial partner BioCat (https://www.biocat.com) or site directed mutagenesis (SDM) can be found in SI appendix Dataset 3. All effector constructs were codon-optimized for *N. benthamiana* using the codon optimization tool of IDT (Integrated DNA technologies, https://eu.idtdna.com/) and synthesized with attL-sites. The constructs were cloned into the binary expression vector pIPKb004 (19) using LR clonase II (Invitrogen) according to the manufacturer and transformed into *A. tumefaciens* using the freeze-thaw protocol described in (20). PCR based SDM and epitope tagging was performed with non-overlapping primers on templates, both listed in SI appendix Dataset 2. Phosphorylation of the linear PCR product was performed with T4 polynucleotide kinase (New England Biolabs) and subsequently ligated with T4 DNA Ligase (New England Biolabs) according to the manufacturer.

#### Section 4 *Agrobacterium tumefaciens* mediated transient expression in *Nicotiana benthamiana*

*Agrobacterium tumefaciens* mediated transient co-expression of effector and resistance genes was conducted according to the protocol described in (7). To test for recognition, effector candidates and resistance genes were infiltrated at a ratio of R:effector 1:4 and incubated for 5 days followed by hypersensitive response quantification by the Fusion Imager FX system as described in (7). Effector genes that did not induce HR response under this standard condition were considered as non-recognised. To allow a quantitative comparison of the recognition strength between *AvrPm17* variants, HR induction was measured in pairwise tests using an infiltration ratio of *Pm17*:*AvrPm17* of 1:1 and infiltrated leaves were imaged at 2dpi or 3dpi depending on the strength of hypersensitive response reaction. HR was subsequently quantified using Fiji (21)) according to the procedure described in (7). Statistical significance was assessed performing a paired wilcoxon rank sum test in Rstudio (v1.2.1335, (11) using the command wilcox.test(paired = TRUE).

#### Section 5 Western blot analysis

Protein extractions were performed according to (7) from eight pooled leaf discs (5mm diameter) originating from four *Agrobacterium* infiltrated leaves at 2dpi. Leaf discs were ground in 150µl 2XLaemmli buffer (100 mM Tris-HCl pH 6.8, 200 mM DTT, 0.04% bromophenol blue, 20% glycerol, 2% SDS), heated to 95°C for 5min, followed by a centrifugation of 10min at 10’000xg. SDS polyacrylamide (PA) gels were used to separate 10 µl of total protein extract (8% PA for PM17-HA and 16% for AVRPM17-FLAG variants), followed by semi-dry blotting on a nitrocellulose membrane (Amersham Protran 0.2 µm NC) using the Trans-Blot SD Semi-Dry Transfer Cell from Bio-Rad. To control for equal loading, the membrane was stained by Ponceau-S. Subsequent detection of HA-tagged proteins was done with a peroxidase-conjugated antibody (anti-HA-HRP, rat monoclonal, clone 3F10, Roche) at a dilution of 1:3000. The FLAG epitope tag was detected with the primary antibody (anti-FLAG, mouse monoclonal, clone M2, Sigma-Aldrich, 1:10’000 dilution). After washing the membrane with TBS-T, the membrane was incubated with anti-mouse peroxidase antibody (anti-mouse-HRP, goat polyclonal, Sigma Aldrich) at a dilution of 1:4000. For chemiluminescence detection, we used the WesternBright ECL HRP substrate (Advansta) and detected the signal with Fusion FX Imaging System (Vilber Lourmat, Eberhardzell, Germany).

**Section 6 Expression analysis**

To verify expression of *AvrPm17* in a selected subset of 16 *B.g tritici* and *B.g. triticale* isolates (Fig. S20) RNA was extracted from leaf segments of the susceptible wheat cultivar ‘Chinese Spring’ 48h after infection with the indicated powdery mildew isolate using the SV total RNA Isolation System Kit (Promega) according to the manufacturer. Uninfected ‘Chinese Spring’ was used as a negative control. Total RNA purity and integrity was verified using a Nanodrop1000 (Thermo Scientific) and agarose gel electrophoresis respectively. Total RNA was reverse transcribed using iScript Advanced cDNA Synthesis Kit for qRT-PCR (Promega) according to the manufacturer. As a negative control, the same reaction was performed in the absence of the reverse transcriptase (‘RTminus’). *AvrPm17*, *AvrPm2* and fungal *GAPDH* were amplified from cDNA using Phusion HF DNA Polymerase (New England Biolabs) and the primers listed in SI appendix Dataset 2. The same reactions were performed on the ‘RTminus’ samples resulting in no detectable PCR product thereby ruling out any contaminations with genomic DNA.

As control, we also performed PCR amplification of *AvrPm17*, *AvrPm2* and fungal *GAPDH* from genomic DNA using the same experimental procedure as described for the cDNA samples above. For extraction of the genomic DNA the susceptible wheat cultivar ‘Chinese Spring’ was infected with the *B.g. tritici* isolate 96224 or the *B.g. triticale* isolate THUN-12 as described above. DNA was extracted 48h after infection using a guanidine-thiocyanate based protocol previously described in (22). Uninfected ‘Chinese Spring’ was used as a negative control. Total genomic DNA purity and integrity was verified using a Nanodrop1000 (Thermo Scientific) and agarose gel electrophoresis respectively. PCR

### Section 7 Bioinformatics analysis

#### Genomic origin of the *AvrPm17* locus

Genomic origin of the *AvrPm17* locus was assessed according to fixed polymorphism in *B.graminis formae specialis* (f.sp*)* following to the rationale described in (13): SNP call was performed on isolates using freebayes (v1.1.0-54-g49413aa, (23)) with the -ploidy 1 setting. Isolates used for this analysis are indicated in SI appendix Dataset 2. The SNP dataset was filtered with vcftools (v0.1.5, (24)). Only SNPs with sequencing depth 20 and quality above 20 were used. Polymorphisms were considered fixed if they were present in all isolates of one f.sp. and absent in all the other ff.spp*.* The polymorphisms found in THUN-12 were compared to this set of fixed polymorphisms. Genomic coverage was calculated using SamTools depth command –a per position and normalized to the average coverage of chromosome 01. Normalized genomic coverage was visualized in 1000bp windows.

#### Structural modelling

Secondary structure prediction was performed using the QUICK2D toolkit (25). Tertiary protein structure modelling was performed using the IntFOLD5.0 server (https://www.reading.ac.uk/bioinf/IntFOLD) (26)

#### Phylogenetic tree

Candidate effectors of the family E003 in *B.g. tritici* 96224 and *B.g. triticale* were identified as described in (2). To identify E003 family members in *B.g. hordei* DH14, the proteins of DH14 were aligned to the E003 effectors using BLASTP+ (v2.6.0, (27)) and hits e-value greater than $e^{-10}$ were retained. Amino acid sequences of the effectors were aligned with MUSCLE (v3.8.31, (28)). The approximativ maximum-likelihood of the phylogenetic relationships were subsequently investigated with the Jones-Taylor-Thorton algorithm implemented with FastTree (v2.1.8, (29)). The local support values of the nearest-neighbor interchanges topology were calculated with the Shimodaira-Hasegawa test from FastTree (v2.1.8,(29). Rooting with the outgroup BGTE-20002 and the graphical representation of the tree were accomplished with FigTree (v1.4.3, http://tree.bio.ed.ac.uk/software/figtree).

#### Cluster analysis

To identify syntenic scaffolds between *B.g. tritici*/*B.g. triticale* and the *B.g.hordei* assembly DH14, we used the Orthofinder tool (v2.3.3, (30)) to identify single orthologous genes in the proteome of *B.g. tritici* 96224 (2) , *B.g. triticale* THUN-12 (unpublished data) and *B.g. hordei* DH14 (31) and *B.g. hordei* RACE1 (2, 31). A scaffold in DH14 was assigned to a chromosome of *B.g. tritici*/*B.g. triticale* if the classification was supported by the majority of the single orthologous genes on the scaffold. Subsequent detailed synteny analysis in the *AvrPm17*-locus was done manually using the integrated genome viewer (IGV, v2.8.6, (16) and genes were considered as syntenic based on conserved orientation and placement in the same orthogroup by OrthoFinder. Low-quality gene models were removed from the analysis if they overlapped with TE, showed no transcriptional support or lacked a start/stop codon. To predict presence/absence of the genes in *B.g. secalis*, Illumina sequences were mapped against the Bgt_genome_v3_16 as described in (2) and copy number variation was estimated using the genomic coverage method described in (2). Therefore, the average number of reads mapping to the genic region of all genes (including intron) were calculated using the SamTools depth –a (v1.7,(32)) command. Average coverage was normalized to the average coverage of all genes. Genes with normalized coverage <0.1 were considered absent. Genes were considered present in the *B.g. secalis* f.sp. if the gene was present in at least one isolate of *B.g. secalis*.

**Haplotype analysis**

Haplotype analysis was conducted based on the re-sequencing data of 160 *B. graminis* isolates as listed in SI appendix Dataset 3. Sequencing of the isolates that were published prior to this study are described in (7, 12, 13). DNA for isolates that were sequenced for this study was extracted using a chloroform/phenol method described in (33) and sequencing was performed as PE150 on the Illumina HiSeq4000 at the functional genomic center Zurich (FCGZ). Copy number of *AvrPm17* genes was estimated using the coverage-based method described above. For the haplotype analysis in the worldwide *B. graminis* population, genomic re-sequencing data was mapped against the Bgt_genome_v3_16 as described in (2). SNP pattern in the genes *Bgt-51729/Bgt-51731* was assessed manually upon visualization in integrative genome viewer IGV (v2.8.6, (16). For isolates where SNP patterns indicate the presence of two distinct *AvrPm17* copies, re-sequencing data were mapped against a single *AvrPm17* gene copy (including 1000bp up and downstream of the gene) and analysed manually with the same approach as described above. Haplovariants from *B.g. dactylidis* were extracted by blasting against previously published de-novo assemblies with BLASTN+ (v2.6.0,(27))

**Gene conversion**

The duplicated regions in the genomes of Bgt_96224 and THUN-12 were aligned using dotter (v4.44.1(34)). The boundaries of the duplicated region were defined manually based on the analysis of the pairwise dot plots, as well as the identification of gene specific insertions. To enable pairwise alignment of the sequences, the insertions identified in Fig. S15 and Fig. S18 were removed. Subsequent pairwise alignment was performed with MUSCLE and nucleotide identity in 50kb sliding windows were calculated using a custom R script. To compare the *AvrPm17* gene of the high quality assemblies of Bgt_96224 and THUN-12 with isolates that carry one gene copy, we performed de-novo assemblies of isolate GZ-6, (see SI appendix Dataset 3) with *SPAdes* (v3.12.0, (35) with the default parameters. BLASTN+ (v2.6.0., (27)) was used to identify the contig containing the *AvrPm17* gene.

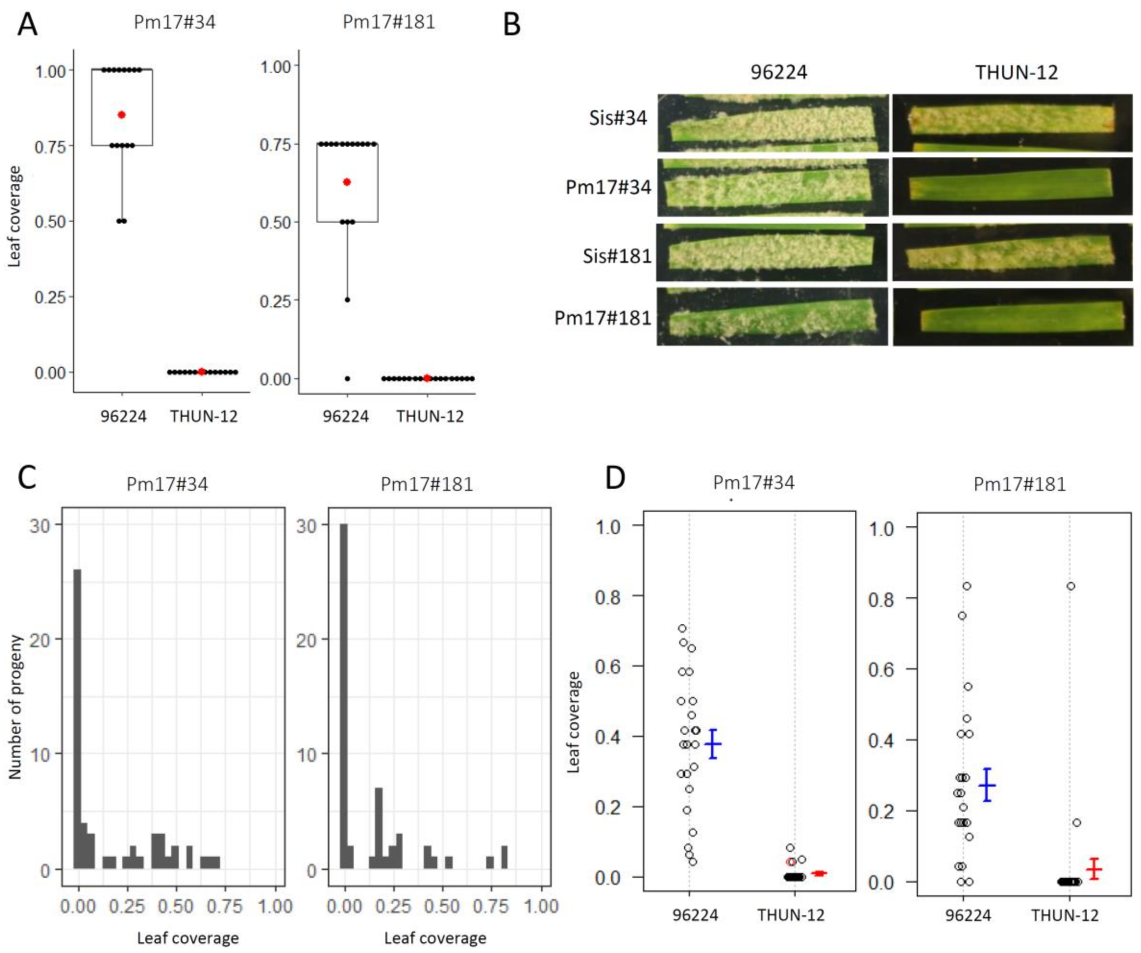

**Fig. S1.** Phenotypes of the mapping population 96224 X THUN-12 on two independent transgenic wheat lines expressing Pm17-HA (A) Phenotype of the parental isolates 96224 and THUN-12 on transgenic lines Pm17#34 and Pm17#181. Leaf coverage of individual leaf segments was scored according to the following scale: avirulent = 0, avirulent/intermediate 0.25, intermediate 0.5, intermediate/virulent =0.75, virulent = 1.  Data from three independent phenotyping experiments are shown. The mean value of all leaf segments is indicated by a red dot (B) representative photographs of phenotypes of 96224 and THUN-12 on the two independent, *Pm17* expressing transgenic lines and their corresponding sister lines at 10dpi (D) Distribution of phenotypes of the 55 randomly selected progeny of the cross 96224 X THUN-12 after 10dpi. (D) phenotypes of the 55 progeny of the cross 96224 X THUN-12 that carry either the 96224 genotype or the THUN-12 genotype at the best associated marker of the QTL.

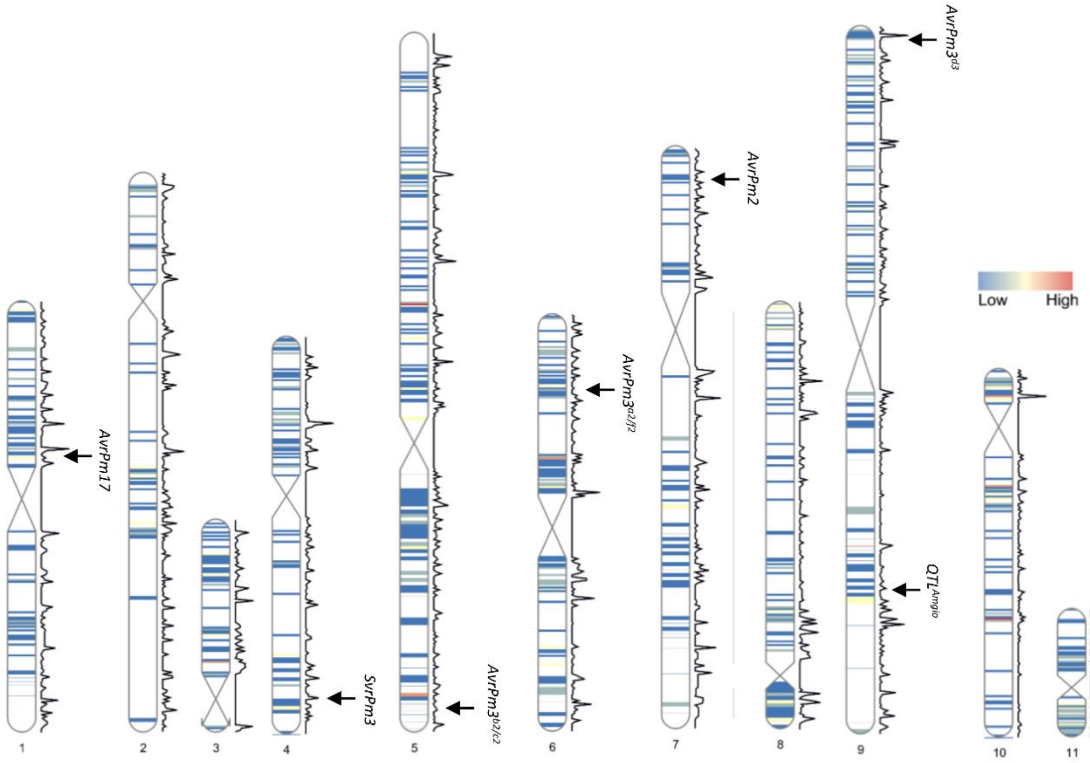

**Fig. S2.** Chromosomal location of *B.g. tritici* avirulence genes and the suppressor of avirulence *SvrPm3* in the assembly of isolate 96224. Chromosomes are colored according to effector gene density in 50kb windows. White areas represents intervals without effectors. Black line represents recombination rate in cM/50kb.

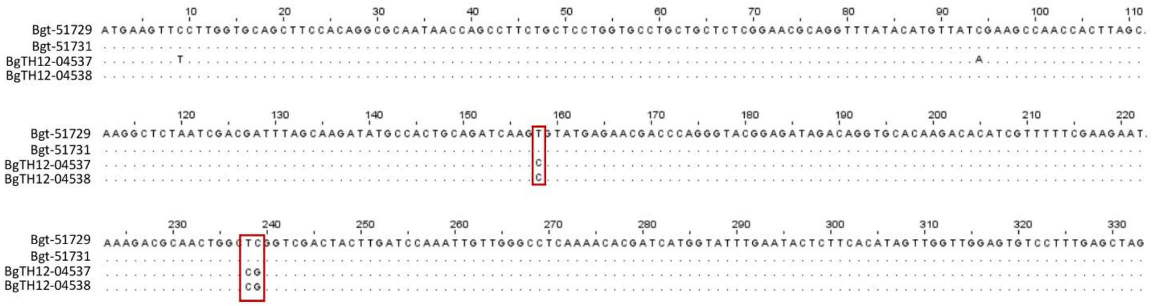

**Fig. S3.** Alignment of the nucleotide sequences of the two genes encoding for AVRPM17 in the isolates Bgt_96224 and THUN-12. Nucleotide differences that lead to amino acid changes are framed with a red box. BgTH12-05637/BgTH12-05438 and Bgt-51729/Bgt-51731 encode for identical proteins.

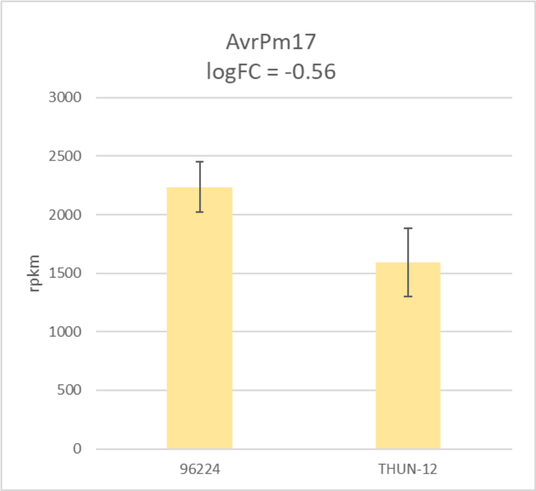

**Fig. S4.** Expression level of *AvrPm17* genes in the two parental isolates Bgt_96224 and THUN-12 using RNAseq data at 2 days post infection. Since *AvrPm17* is encoded by identical gene copies in the isolate 96224 and almost identical copies in THUN-12, expression levels can only be quantified combined for both gene copies. Expression levels are indicated as rpkm (reads per kilobase per million reads), logFC (log-fold changes) of >1.5 are considered significant.

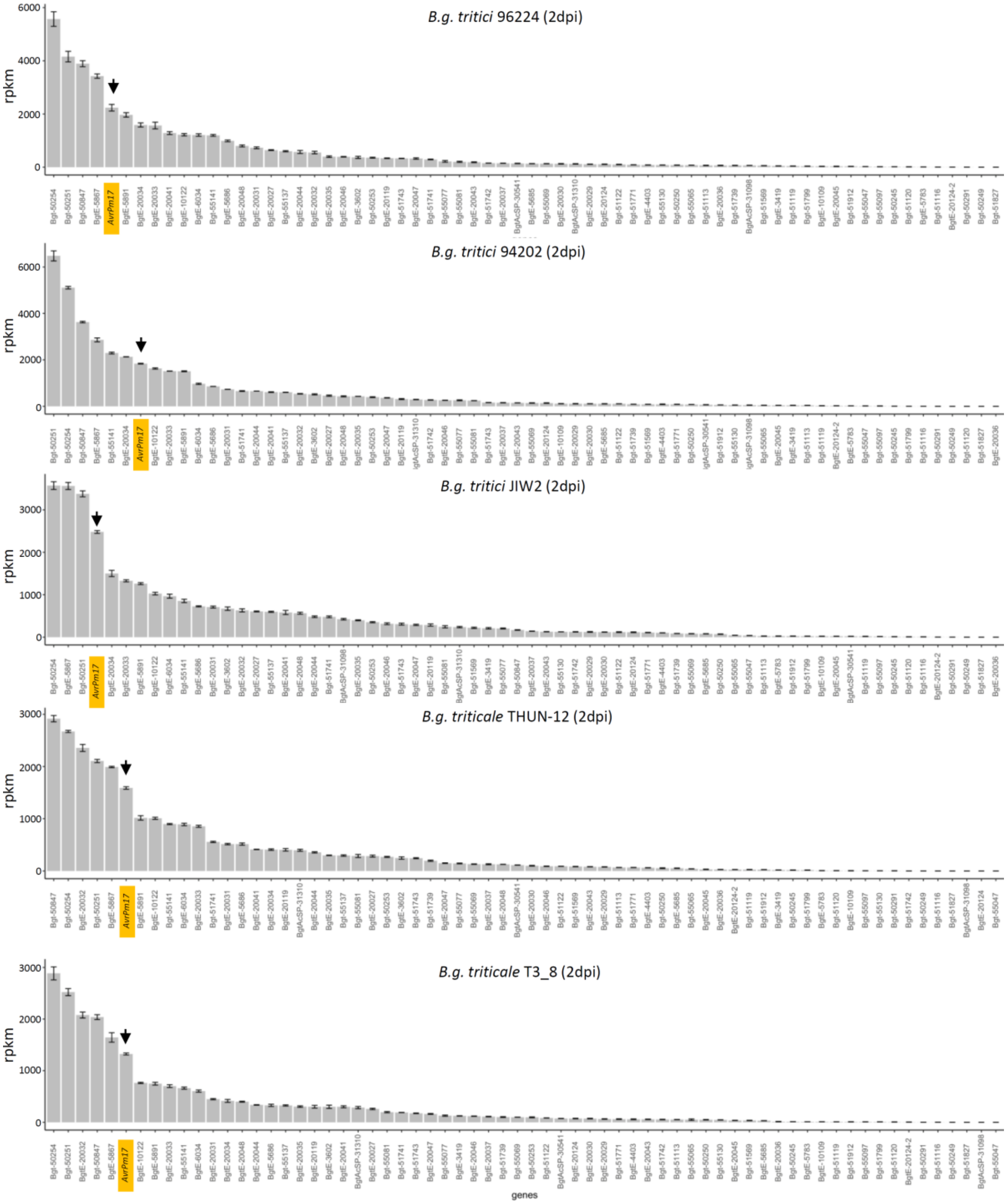

**Fig. S5***.* Expression of genes encoding members of the candidate effector family E003 in three *B.g. tritici* isolates and two *B.g. triticale* isolates at 2dpi (days post infection) using RNAseq data. Expression levels are indicated as rpkm (reads per kilobase per million reads) and represent the average of three biological replicates for each isolate. *AvrPm17* (*Bgt-51729/Bgt-51731*) is highlighted in yellow.

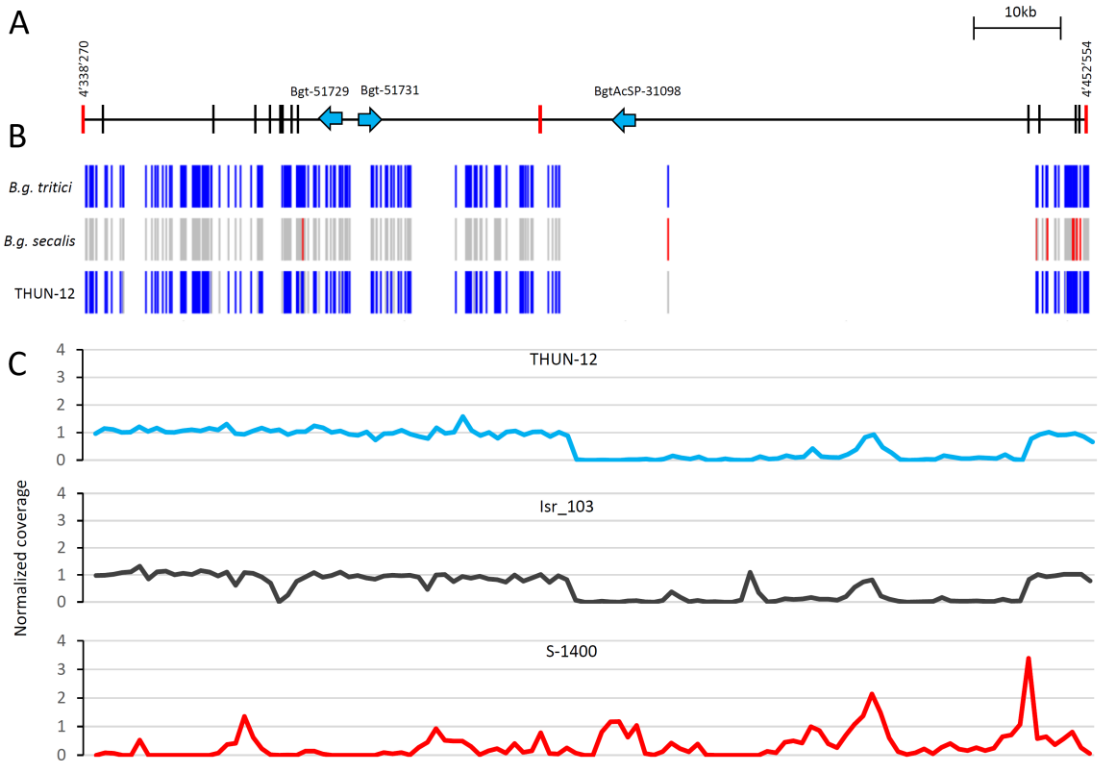

**Fig. S6.** Genomic origin of the *AvrPm17* locus on chromosome 01 in *B.g. triticale* isolate THUN-12. (A) representation of the physical interval of the *AvrPm17* locus in the Bgt_96224 genome assembly. Gene size is not drawn to scale. Bars represent informative markers in the genetic confidence interval (1.5LOD). Flanking markers and the best associated marker of the QTL are depicted in red.. (B) Determination of genomic origin based on fixed polymorphisms within *formae speciales*. Blue bars represent polymorphisms fixed in *B.g. tritici* that differ from *B.g. secalis*, red lines represent polymorphisms that are fixed in *B.g. secalis* and differ from *B.g. tritici.* Grey indicates SNPs that could not be determined in *B.g. secalis* or THUN-12 because of lacking genomic coverage likely due to deletions of these particular regions. (C) Genomic coverage of the selected isolates *B.g. triticale* THUN-12, *B.g. tritici* Isr_103 and *B.g. secalis* S-1400. The 50kb deletion identified in *B.g. triticale* isolate THUN-12 can also be found in certain *B.g. tritici* isolates such as Isr_103. Normalized genomic coverage was estimated in 1000bp windows normalized to the average coverage of Bgt_chr-01.

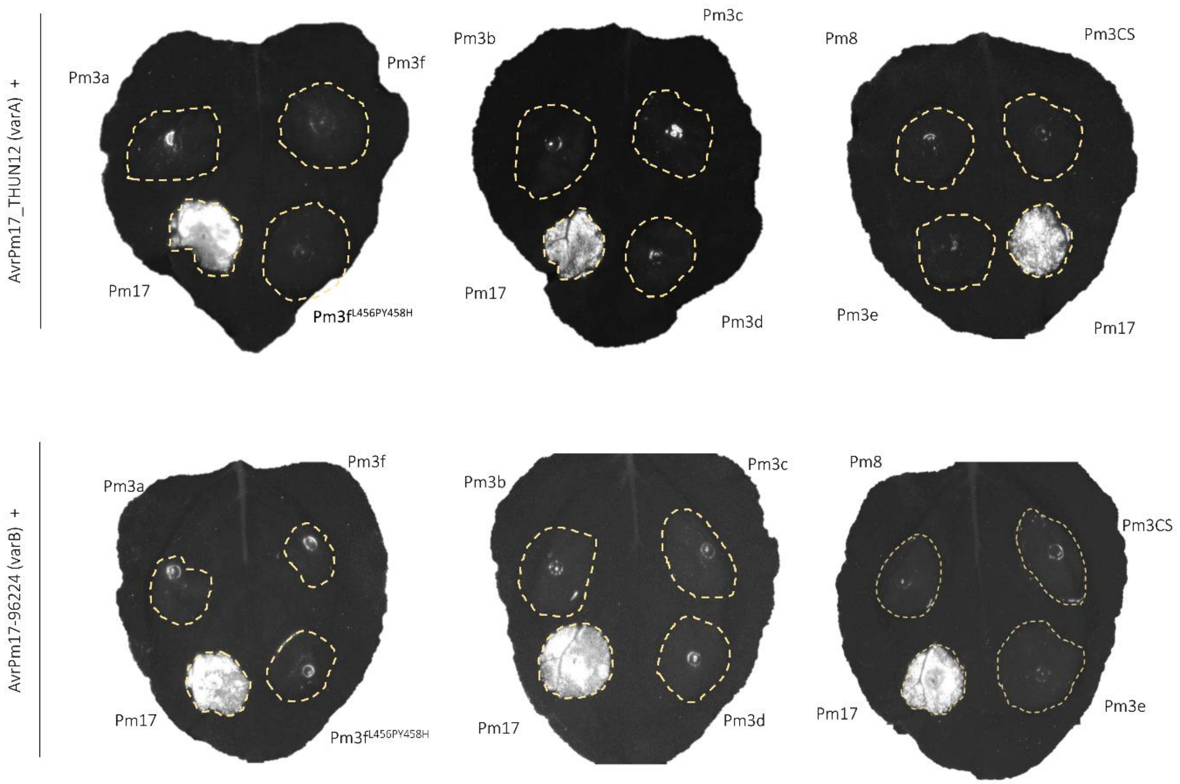

**Fig. S7.** AVRPM17 is not recognized by other members of the Pm3-NLR family. *AvrPm17_THUN12* (varA) and *AvrPm17_96224* (varB) were co-expressed with different alleles of the *Pm3/8/17* family. All infiltrations were done with the ratio R: Avr of 1:4 and imaged after 5dpi. Each infiltration was done with at least n=4 leaves and repeated twice.

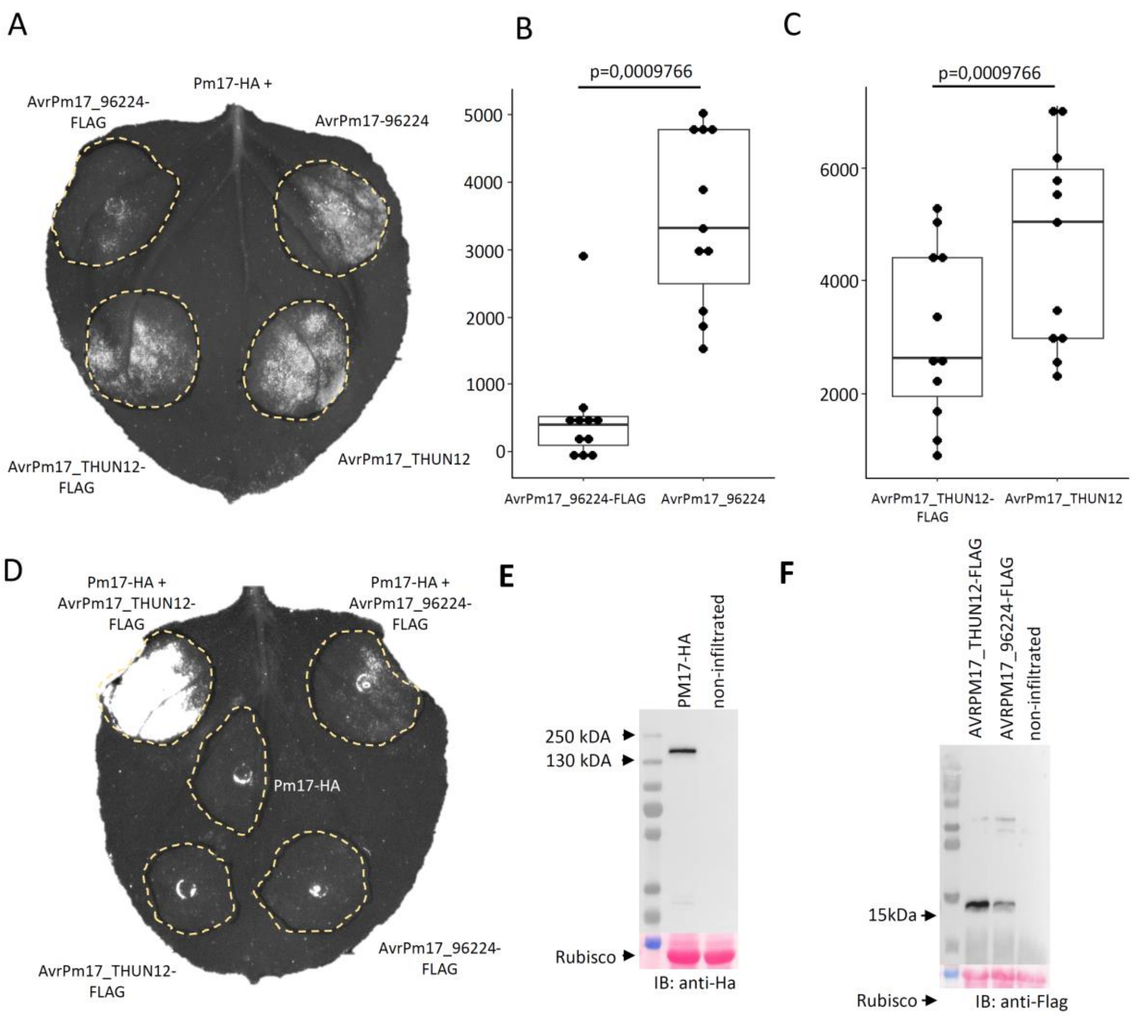

**Fig. S8.** Epitope tagging and western blot analysis of PM17 and AVRPM17 variants. (A-C) C-terminal FLAG-tag interferes with HR induction of AVRPM17 upon co-infiltration with PM17-HA. (A) shows a representative agrobacterium-infiltrated *N. benthamiana* leaf expressing Pm17-HA with FLAG epitope tagged and untagged versions. Pm17-HA and AVRPM17 variants were infiltrated at 1:4 ratios. (B,C) HR of n=10 leaves was quantified at 2dpi in two independent experiments. Statistical significance was accessed using the paired Wilcoxon ranked sum test, p-values are reported above the boxplots. (D) shows a representative picture of a agrobacterium-infiltrated leaf expressing PM17-HA with AVRPM17-FLAG variants at a R:AVR ratio of 1:6 and individual infiltrated constructs. HR of n=6 leaves was measured after 5dpi with the Fusion ImagerX system in two different experiments (E,F) western blot of Pm17-HA and AVRPM17-FLAG variants.

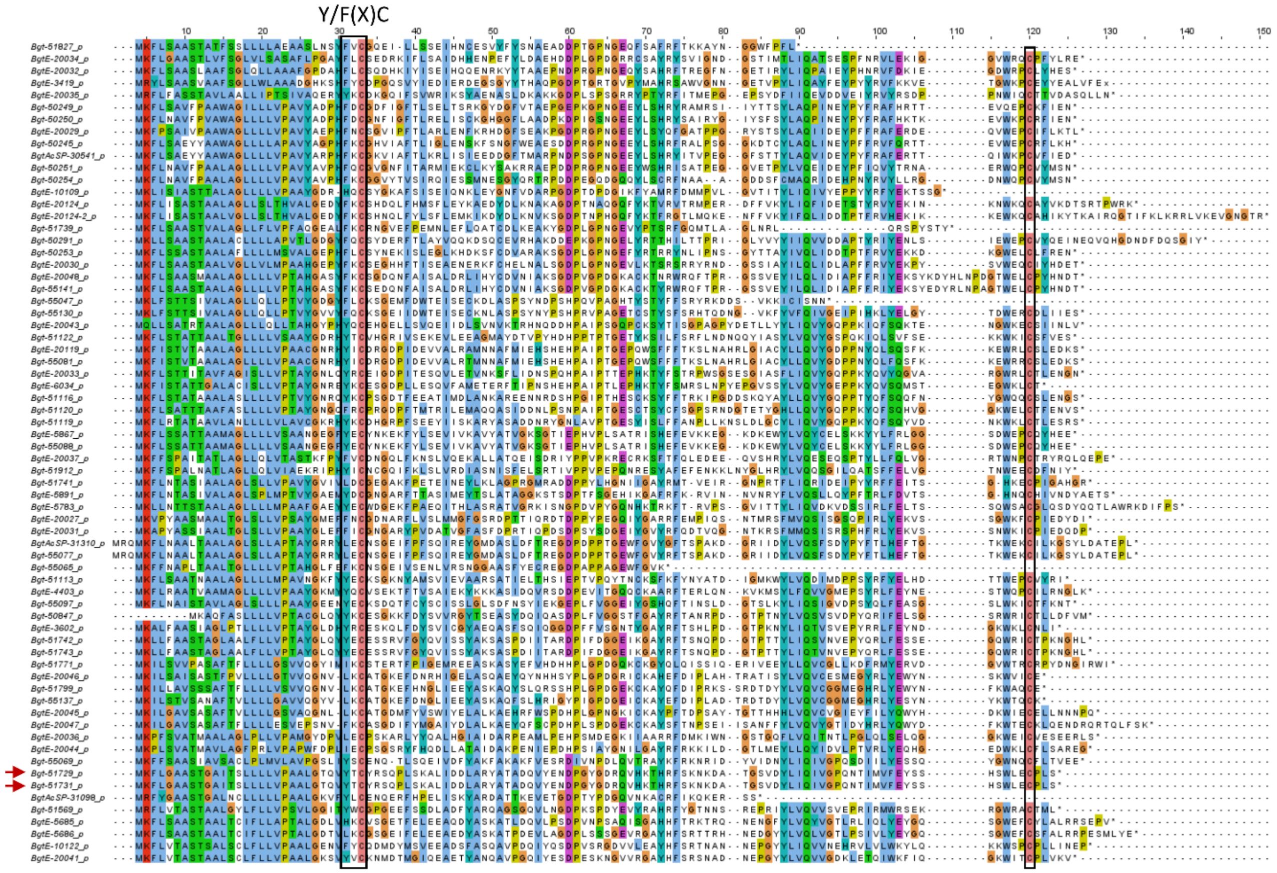

**Fig. S9**. Protein sequence alignments of the effector family E003 members found in the reference isolate Bgt_96224. AVRPM17 is highlighted by red arrows. The conserved Y/F(x)C motif and C-terminal cysteine are indicated (black boxes). The alignment was colored according to the ClustalX color scheme as follows: blue: hydrophobic; red: positively charged; purple: negatively charged; green: polar uncharged; yellow: proline; fleshy pink: cysteine; orange: glycine.

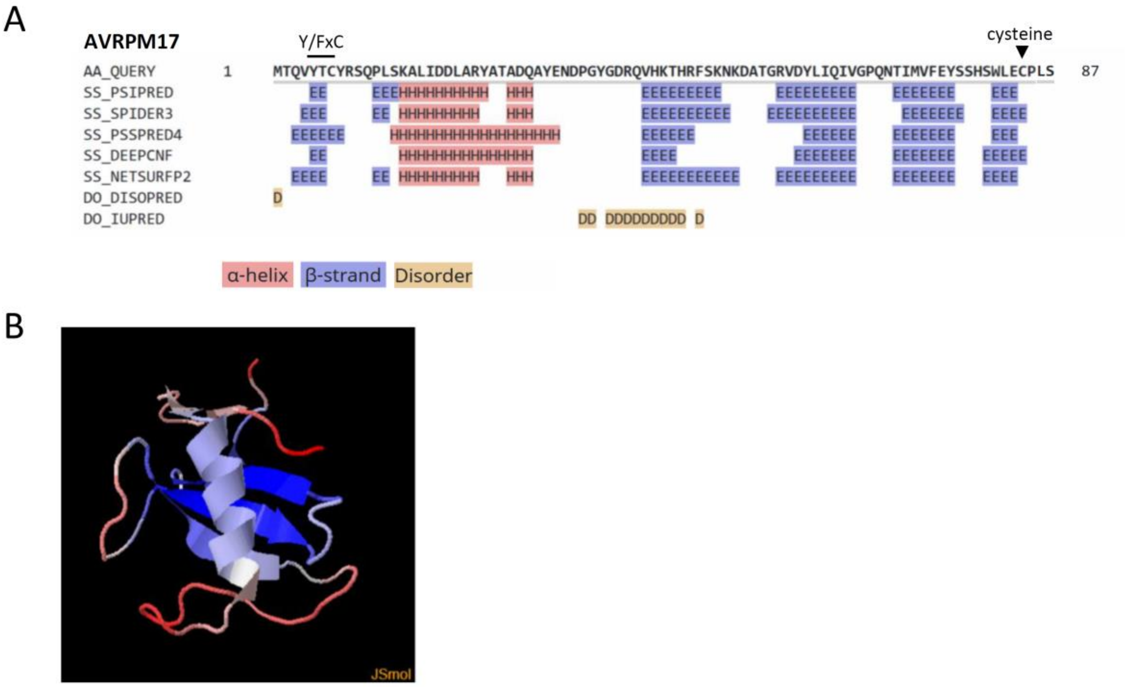

**Fig. S10**. Structural modelling of the AVRPM17 protein. (A) Modelling of the 2D protein structure of AVRPM17_THUN12 without signal peptide using the the Quick2D toolkit**.** Y/FxC motif and C-terminal cysteine are indicated (B) Representation of the best model (p value=1.145E-4) obtained by modelling the mature peptide of AVRPM17_THUN-12 with IntFOLD5.0. The model is based on the following templates: fungal RNAse po1 from *Pleurotus ostreatus* (3whoA, described in (36)) and BEC1054 an effector from *B.g. hordei* (6fmbA, described in (37)) The model is colored according to the JSmol coloring scheme in which blue is designating high model accuracy and red low model accuracy.

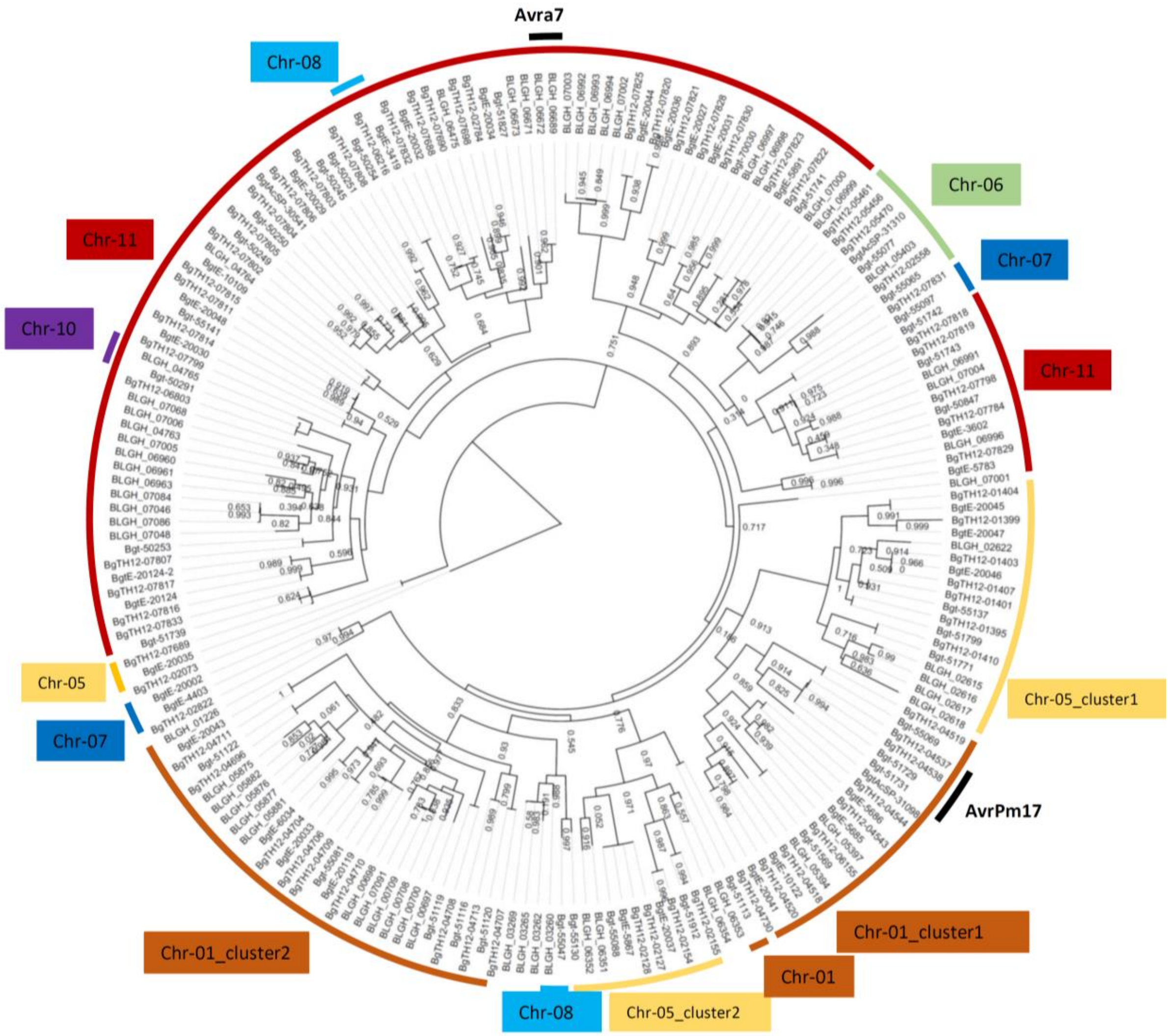

**Fig.S11.** Phylogenetic tree of the E003 family with 69 members in *B.g. tritici*, 70 members in *B.g. triticale* THUN-12 and 59 members in DH14. Gene clades were colored according to their position on the eleven chromosomes of *B.g tritici* and *B.g. triticale*. The local support values to the nearest-neighbor interchanges topology were calculated with the Shimodaira-Hasegawa test and indicated for each branch.

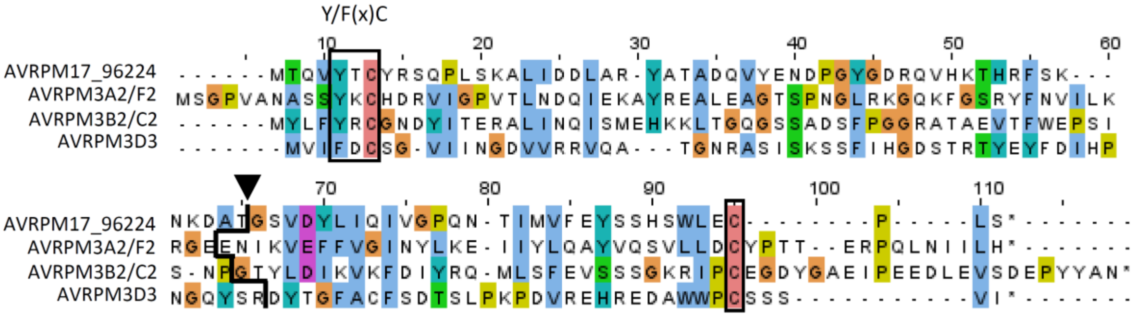

**Fig. S12.** Protein alignment of the mature protein (without signal peptide) of AVRPM17_96224, AVRPM3$^{A2/F2}$, AVRPM3$^{B2/C2}$ and AVRPM3$^{D3}$. The Y/F(x)C motif and the conserved cysteine are indicated with black boxes. The intron position in AVRPM17 is indicated by a black arrow and by a black line in the AVRPM3s. The alignment was colored according to the ClustalX color scheme as follows: blue: hydrophobic; red: positively charged; purple: negatively charged; green: polar uncharged; yellow: proline; pink: cysteine; orange: glycine.

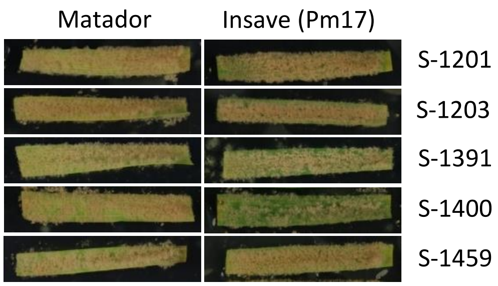

**Fig. S13**. Phenotypes of five *B.g. secalis* isolate on the rye *Pm17* donor ‘Insave’ and the susceptible rye cultivar Matador. Images were taken at 10dpi.

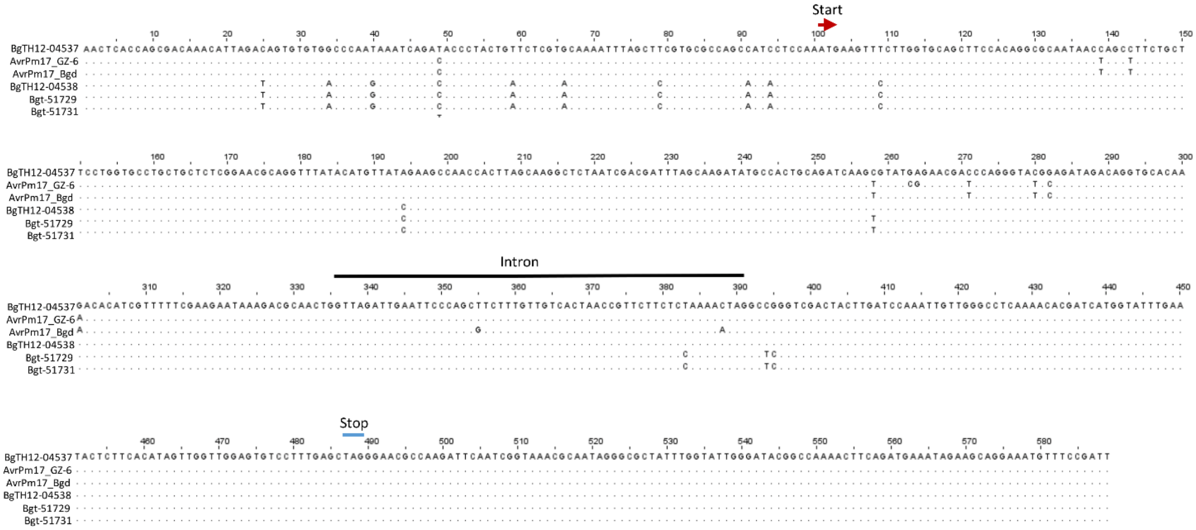

**Fig. S14.** Alignment of nucleotide sequences of *AvrPm17* in different *B. graminis* isolates. Alignment encompasses the coding sequences and 100bp up- and downstream of the start and stop codon respectively. Position of start and stop codons and the intron are indicated

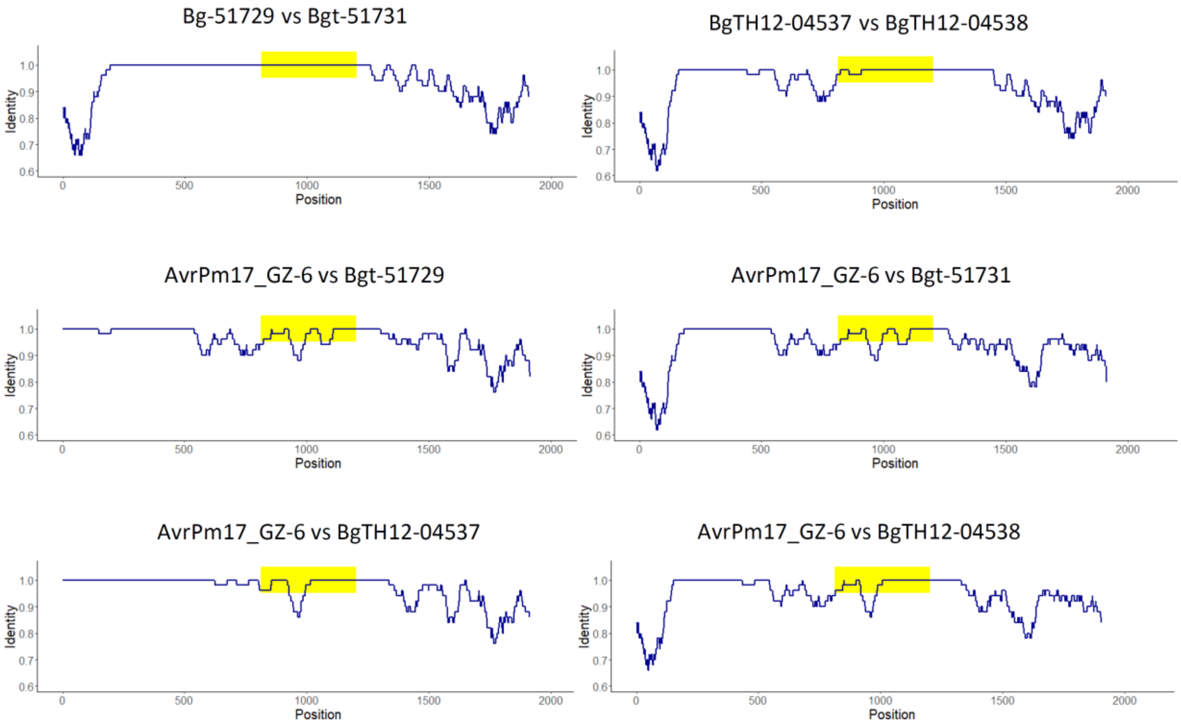

**Fig. S15** Visual representation of sequence alignments of the *AvrPm17* gene duplication in different isolates. The x-axis shows the alignment position, while the y-axis shows the sequence identity calculated in 50bp sliding window. To facilitate alignment, the insertions/deletions in the sequences (see Fig. S15, Fig. S18) were removed. The position of the *AvrPm17* gene is highlighted in yellow. Note that in the alignment of *Bgt-51729* and *Bgt-51731* (first panel), the region containing the gene shows 100% identity while sequence identity decreases toward the end of the duplicated regions. The 200bp flanking region show 79% up and 84% downstream respectively. This suggests recurring gene conversions that keep the two copies of the gene identical.

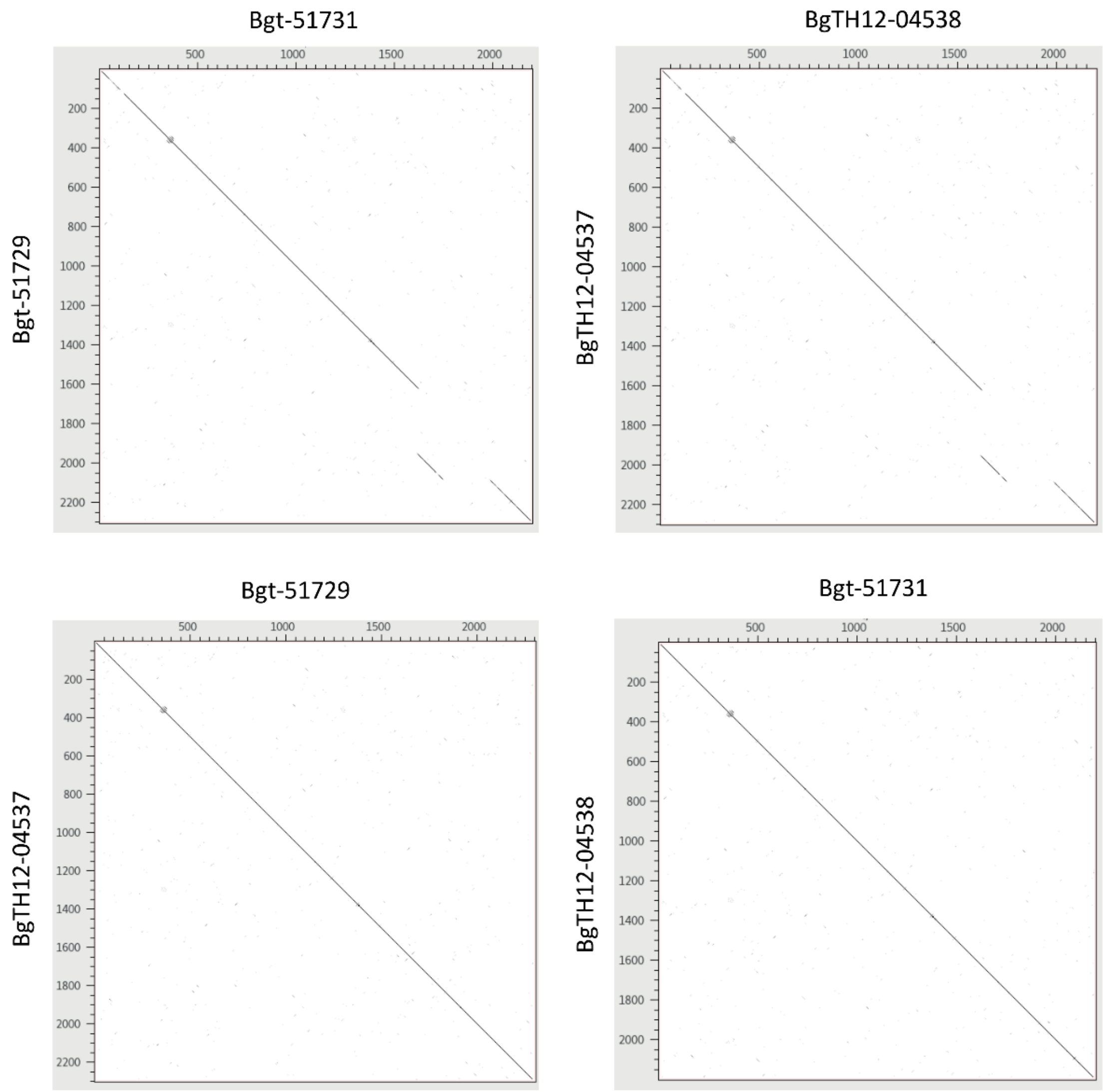

**Fig. S16.** Pairwise dotplots of the duplicated *AvrPm17* genes and flanking regions in the high-quality genomes of Bgt_96224 and THUN-12. The duplicated region spans 813 upstream of the start codons of the four genes and 1,079 and 998 downstream of the stop codon of *Bgt-51729/BgTH12-04537* and *Bgt-51731/BgTH12-04538*, respectively. The duplicated regions spanning 2,300bp for *Bgt-51729/BgTH12-04537* and 2200bp for *Bgt-51731/BgTH12-04538* were aligned using dotter.

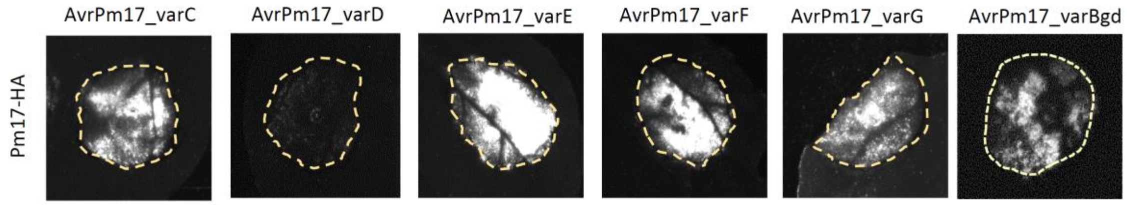

**Fig. S17.** Co-expression of *AvrPm17* haplovariants with *Pm17*-HA in *Nicotiana benthamiana*. Infiltrations were performed at infiltration ratio R:AVR of 1:4 and HR development was imaged using the Fusion Imager FX system after 5dpi. Effectors were considered as recognized if they induced an HR response under these standard conditions.

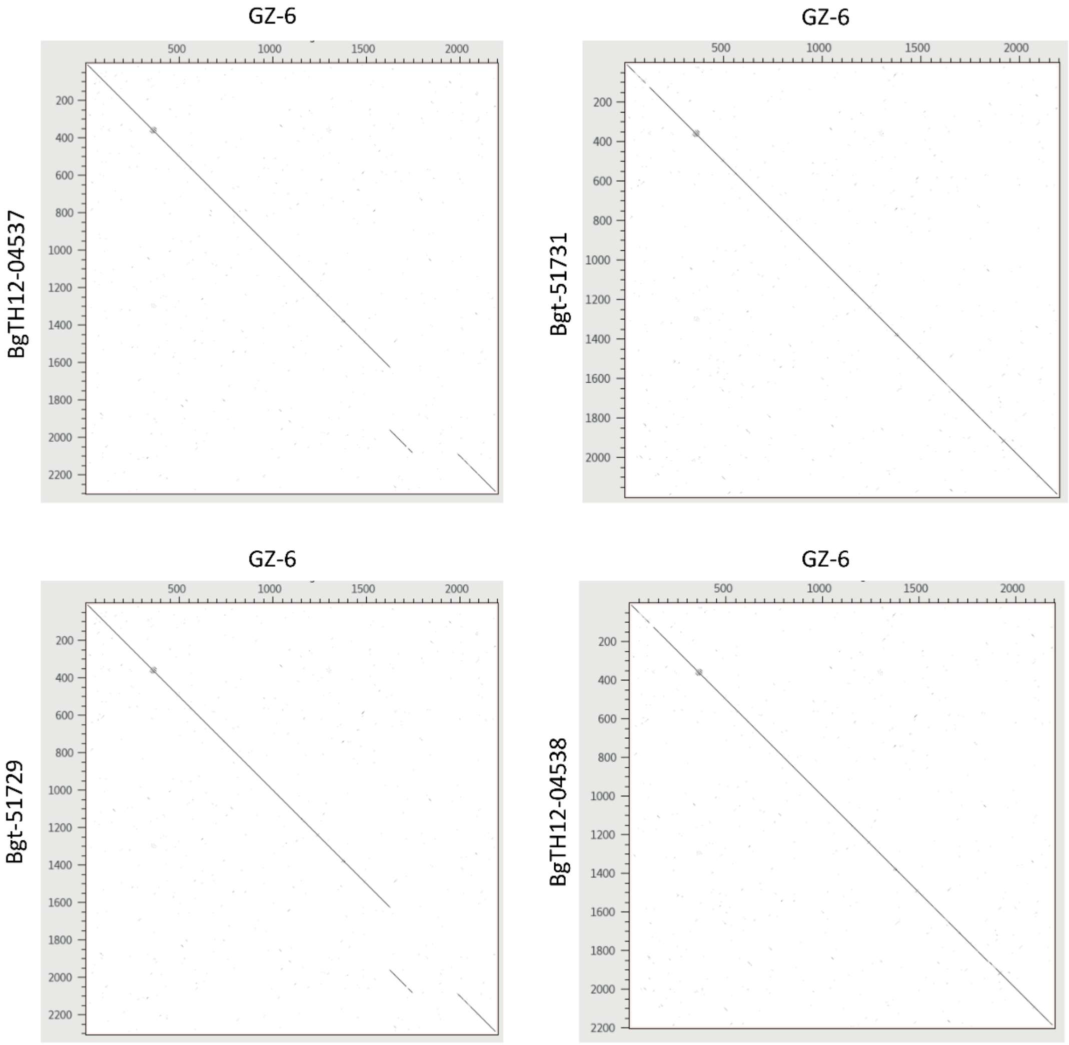

**Fig. S18.** Pairwise dotplots of the duplicated *AvrPm17* genes and flanking region in the high-quality genome of Bgt_96224 and THUN-12 with the *AvrPm17* gene copy of isolate GZ-6**.** Isolate GZ-6 contains only one *AvrPm17* copy. The upstream flanking region of this gene is more similar to *BgTH12-04537/Bgt-51729* as shown in the left panels, whereas the downstream region of the gene in GZ-6 is more similar to *BgTH12-04538/Bgt-51731* as shown in the right panels. This indicates that the *AvrPm17* gene copy is a recombinant between the two duplicated gene copies.

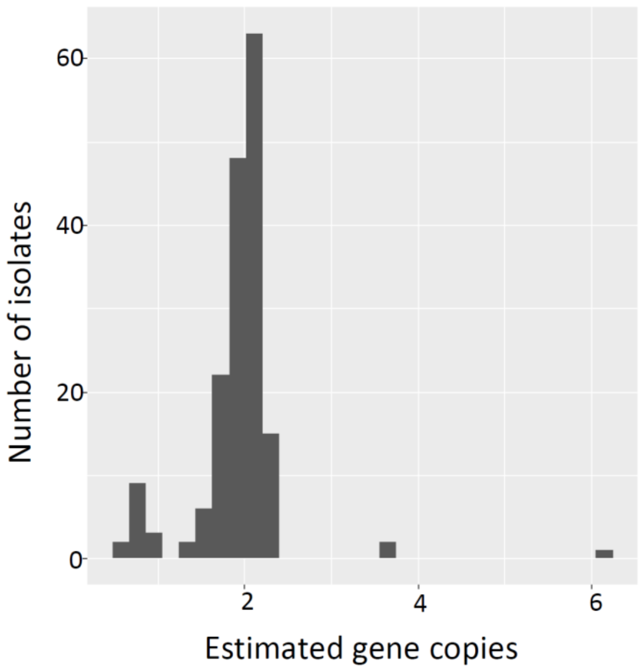

**Fig. S19** The majority of *B.g. tritici*, *B.g. triticale* and *B.g. dicocci* isolates encode for two copies of *AvrPm17*. Copy number variation was estimated using genomic coverage of resequencing data normalized by the genomic coverage of all genes.

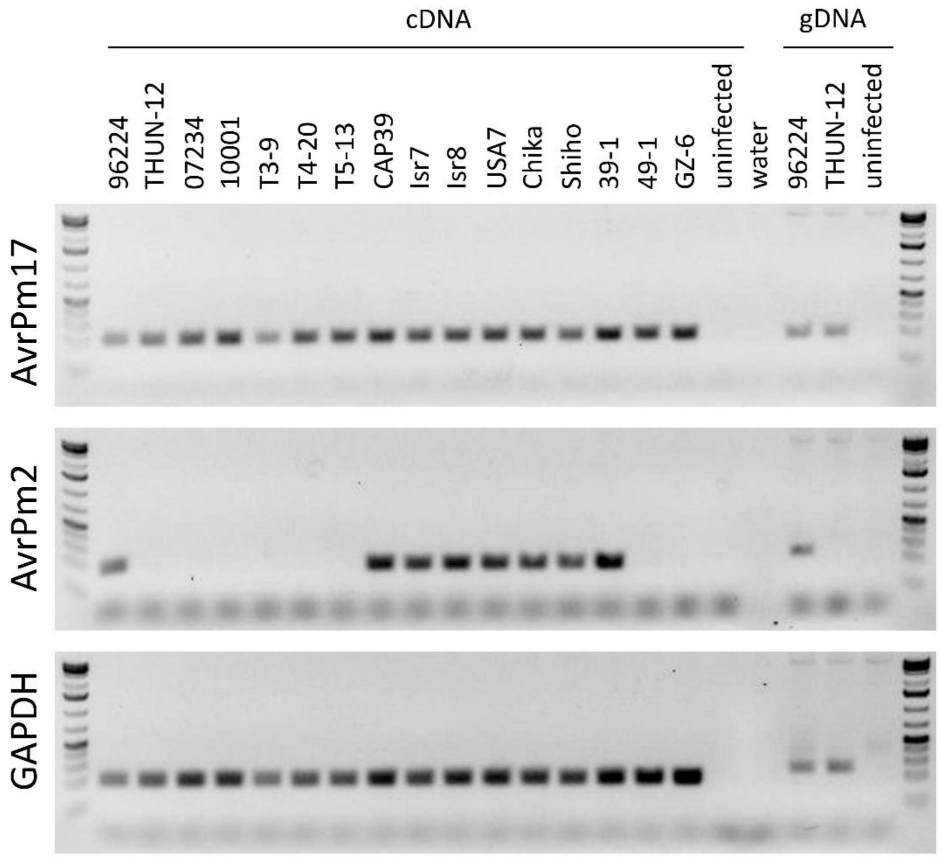

**Fig. S20**. *AvrPm17* is expressed in all tested isolates at 2dpi. Expression of the *AvrPm17* genes was tested using RT-PCR on cDNA originating from infected leaf segments of the susceptible cultivar Chinese Spring. Uninfected Chinese Spring was used as a control. RT-PCR amplification of the fungal housekeeping gene GAPDH and *AvrPm2*, which displays a well-characterized presence/absence polymorphism (12) are shown for comparison. PCR amplification of *AvrPm17, AvrPm2* and fungal *GAPDH* from genomic DNA of the two reference isolates 96224 and THUN-12 or uninfected Chinese Spring are shown for comparison. Amplicon size is slightly bigger on gDNA due to amplification over a small intron for all three tested genes.

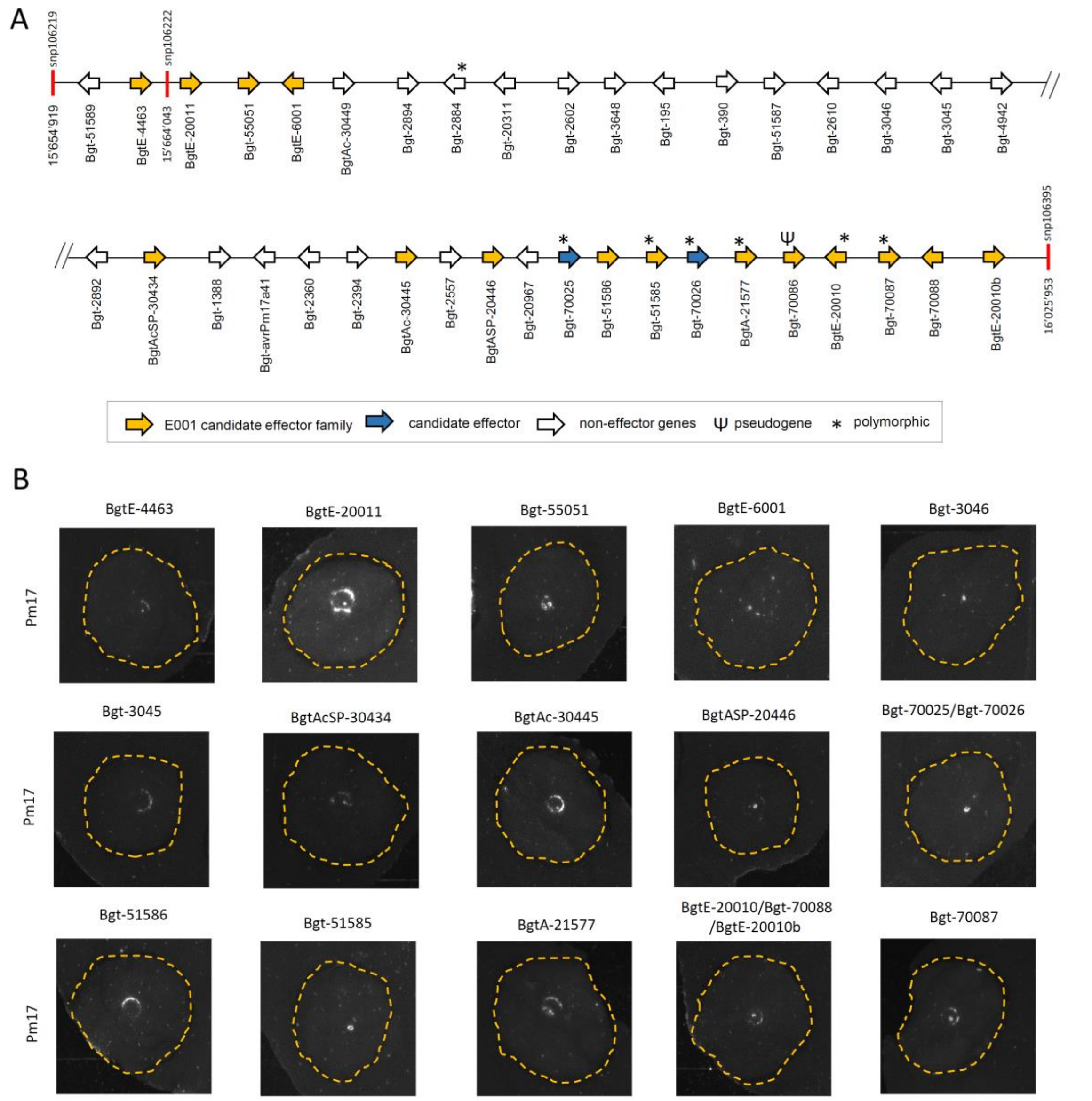

**Fig. S21.** QTL on chromosome 9 contains numerous effector candidates but does not encode a *Pm17* avirulence component. (A) Schematic representation of the physical interval underlying the genetic confidence interval (LOD=1.5) on chromosome 9 on the assembly of the avirulent isolate 96224. Genes and orientation are indicated; gene size is not drawn to scale. Polymorphic genes between parental isolates 96224 and THUN-12 are indicated by an asterisk. Bgt-70025 and Bgt-70026 represent genes that contain a signal peptide and show no homology outside the genus *Blumeria* and therefore likely represent candidate effectors. (B) *N. benthamiana* co-expression assays with Pm17-HA of the 16 effector genes and two putatively secreted non-effector genes found within the confidence interval depicted in panel (A). Infiltrations were performed at a ratio of Pm17: candidate gene of 1:4 and images were taken 5 days after infiltration using the FX Fusion Imager system.

**Table S1.** Results of the QTL mapping

| Line | Chr | Marker[a] | cM[b] | LOD[c] | Pos[d] | 1.5LOD[e] | Interval Bgt_96224[f] | Interval THUN-12[g] |
|---|---|---|---|---|---|---|---|---|
| BW Pm17#34 | Bgt_chr-01 | snp4585 | 164.8 | 9.2 | 4390396 | snp4525 -snp4614 | 4'338'270 -4'452'554 | 4'085'291 -4'147'189 |
| BW Pm17#181 | Bgt_chr-01 | snp4552 -snp4584 | 164.8 | 7.0 | 4361087 -4390173 | snp4394 -snp4622 | 4'166'104 -4'458'811 | 3'921'779 -4'159'203 |

[a]Best associated marker in the single interval QTL analysis. It is possible that several markers are equally significant

[b]cM position at the best associated marker

[c]Logarithm of the odds. Significance LOD threshold was calculated by 1000 permutations

[d]Position of the best associated SNP in the Bgt_genome_v3_16 assembly

[e]Markers delimitating the genetic confidence interval (1.5LOD interval)

[f]Physical interval underlying the genetic confidence interval in the *B.g. tritici* assembly Bgt_genome_v3_16

[g]Physical interval underlying the genetic confidence interval in the *B.g. triticale* assembly THUN12_genome_v1

**Table S2.** Nucleotide identity in the 200bp distal region of the gene duplication in different isolates. Upper panel represents value of the upstream region of the duplication, lower panel represent the region downstream of the genes.

| | Bgt-51729 | BgTH12-04537 | AvrPm17_GZ-6 | BgTH12-04538 | Bgt-51731 |
|---|---|---|---|---|---|
| Bgt-51729 | 100 | 99.5 | 99.5 | 78 | 79 |
| BgTH12-04537 | 99.5 | 100 | 100 | 78.5 | 79.5 |
| AvrPm17_GZ-6 | 99.5 | 100 | 100 | 78.5 | 79.5 |
| BgTH12-04538 | 78 | 78.5 | 78.5 | 100 | 99 |
| Bgt-51731 | 79 | 79.5 | 79.5 | 99 | 100 |

| | Bgt-51729 | BgTH12-04537 | Bgt-51731 | BgTH12-04538 | AvrPm17_GZ-6 |
|---|---|---|---|---|---|
| Bgt-51729 | 100 | 99.49 | 84 | 84 | 86.5 |
| BgTH12-04537 | 99.49 | 100 | 83.33 | 83.33 | 85.86 |
| Bgt-51731 | 84 | 83.33 | 100 | 100 | 90 |
| BgTH12-04538 | 84 | 83.33 | 100 | 100 | 90 |
| AvrPm17_GZ-6 | 86.5 | 85.86 | 90 | 90 | 100 |

**Table S3.** Phenotypes of natural *B.g. tritici* and *B.g. triticale* isolates on *Pm17* transgenic lines.

| *f.sp.* | Haplotype[a] | Pm17#34[b] | Pm17#181[b] | Isolate |
|---|---|---|---|---|
| *B.g. tritici* | varA/varA | 0.00 | 0.00 | 97223 |
| *B.g. triticale* | varA/varA | NA | 0.00 | T3-9 |
| *B.g. tritici* | varA/varA | 1.00 | 1.00 | 39-1 |
| *B.g. tritici* | varA/varC | 0.46 | 0.17 | 10001 |
| *B.g. tritici* | varB/varB | 0.90 | 0.92 | 21-1 |
| *B.g. tritici* | varB/varB | 0.71 | 0.54 | Isr70 |
| *B.g. tritici* | varB/varB | 0.33 | 0.25 | 85063 |
| *B.g. tritici* | varB/varB | 0.63 | 0.58 | Usa7 |
| *B.g. tritici* | varC/varC | 0.33 | 0.25 | Isr7 |
| *B.g. triticale* | varC/varC | 0.83 | 0.29 | T5-13 |
| *B.g. tritici* | varC/varC | 0.83 | 0.42 | Isr8 |
| *B.g. tritici* | varC/varC | 0.50 | 0.25 | Shiho |
| *B.g. triticale* | varC/varC | 0.33 | 0.00 | CAP-39 |
| *B.g. tritici* | varC/varC | 0.38 | 0.44 | Chikara |
| *B.g. tritici* | varD | 1.00 | 0.96 | GZ-6 |
| *B.g. tritici* | varD | 0.96 | 0.96 | 49-1 |

[a] AVRPM17 haplotype combination encoded by the isolate
**[b]** Leaf coverage of individual leaf segments was scored according to the following scale: avirulent = 0, avirulent/intermediate 0.25, intermediate 0.5, intermediate/virulent =0.75, virulent = 1. For each isolate, the average of at least 4 individual leaf segments is indicated.

**Table S4. Results of the QTL mapping based on 117 progeny of the cross Bgt_96224 X THUN12**

| Line | Chr | Marker[a] | cM[b] | LOD[c] | Pos[d] | 1.5LOD[e] | Interval 96224[f] |
|---|---|---|---|---|---|---|---|
| Amigo | Bgt_chr-09 | c9.loc340 | 340.0 | 15.97 | 15’664’043 | snp106219<br>snp106395 | 15’654’919<br>16’025’953 |
| Amigo | Bgt_chr-01 | snp4594<br>-snp4610 | 165.8 | 4.24 | 4’357’972 | snp4389<br>snp4685 | 4’157’551<br>4’530’020 |

[a]Best associated Marker in the single interval QTL analysis
[b]cM position of the best associated marker
[c]Logarithm of the odds, significance LOD threshold was calculated by 1000 permutations
[d]Position of best associated SNP in the *B.g. tritici* assembly Bgt_genome_v3_16
[e]Markers delimitating the genetic confidence interval (1.5LOD interval)
[f]Physical interval underlying the genetic confidence interval in the *B.g. tritici* assembly Bgt_genome_v3_16

**Table S5.** ***Summary of genes in genetic confidence interval on chromosome 9***

| Name | SP[a] | Polymorphic[b] | Gene description[c] | Effector family[d] | logFC[e] | Expression 96224[f] | Expression THUN-12[g] | Tested[h] |
|---|---|---|---|---|---|---|---|---|
| Bgt-51589 | - | - | - | - | - | 0.0 | 0.0 | |
| BgtE-4463 | - | - | Candidate effector | E001 | 0.00 | 19.4 | 19.4 | yes |
| BgtE-20011 | - | - | Candidate effector | E001 | -0.09 | 47.3 | 46.5 | yes |
| Bgt-55051 | yes | - | Candidate effector | E001 | -0.49 | 37.0 | 27.1 | yes |
| BgtE-6001 | yes | - | Candidate effector | - | -0.36 | 52.7 | 41.7 | yes |
| BgtAc-30449 | - | - | - | - | -0.41 | 51.4 | 40.6 | |
| Bgt-2894 | - | - | Putative ntf2-like domain containing protein | - | 0.08 | 124.6 | 140.7 | |
| Bgt-2884 | - | 1 | DUF962-domain containing protein | - | 0.09 | 141.0 | 155.0 | |
| Bgt-20311 | - | - | RIX_Bgt_Tara | - | -0.62 | 55.7 | 38.0 | |
| Bgt-2602 | - | - | GTPase-activating protein | - | 0.36 | 16.6 | 23.3 | |
| Bgt-3648 | - | - | - | - | 0.63 | 11.6 | 18.4 | |
| Bgt-195 | - | - | O-mannosyltransferase | - | -0.04 | 103.2 | 105.0 | |
| Bgt-390 | - | - | Dipeptidyl-aminopeptidase B | - | 0.11 | 42.1 | 48.5 | |
| Bgt-51587 | - | - | - | - | - | 0.0 | 0.0 | |
| Bgt-2610 | - | - | Ribokinase-like protein | - | 0.13 | 122.3 | 141.4 | |
| Bgt-3046 | yes | - | putative upf0480 protein | - | -0.46 | 110.9 | 83.3 | yes |
| Bgt-3045 | yes | - | MARVEL-like domain protein | - | -0.88 | 58.8 | 33.3 | yes |
| Bgt-4942 | - | - | O-acetylhomoserine (thiol)-lyase | - | -0.16 | 41.1 | 39.1 | |
| Bgt-2892 | - | - | RNA polymerase II subunit B16 | - | -0.51 | 209.1 | 157.1 | |
| BgtAcSP-30434 | yes | - | Candidate effector | E001 | - | 2.2 | 0.7 | yes |
| Bgt-1388 | - | - | Ergosterol biosynthesis | - | -1.18 | 299.2 | 139.3 | |
| Bgt-avrPm17a41 | - | - | Putative RNA-binding protein fus tls protein | - | 0.02 | 99.8 | 107.0 | |
| Bgt-2390 | - | - | Rab2/secretion related GTPase | - | -0.12 | 37.3 | 36.0 | |
| Bgt-2394 | - | - | Choline transporter-like protein | - | 0.29 | 42.3 | 54.0 | |
| BgtAc-30445 | - | - | Candidate effector | - | -0.14 | 12.6 | 11.7 | yes |
| Bgt-2557 | - | - | Transcription factor | - | -0.03 | 31.4 | 33.8 | |
| BgtASP-20446 | yes | - | Candidate effector | E001 | -0.40 | 48.0 | 38.2 | yes |
| Bgt-20967 | - | - | No open reading frame | - | - | 21.8 | 18.4 | |
| Bgt-70025 | yes | 2 | Candidate effector | NC | 0.60 | 39.7 | 67.0 | yes |

| | | | | | | | | |
|---|---|---|---|---|---|---|---|---|
| Bgt-51586 | yes | - | Candidate effector | E001 | 0.31 | 18.9 | 24.7 | yes |
| Bgt-51585 | yes | 2 | Candidate effector | E001 | -0.11 | 56.6 | 54.9 | yes |
| Bgt-70026 | yes | 1 | Candidate effector | NC | -0.86 | 21.7 | 12.3 | yes |
| BgtA-21577 | - | 4,0[i] | Candidate effector | E001 | 0.63 | 22.9 | 36.3 | yes |
| Bgt-70086 | - | - | Candidate effector, premature stop codon | - | -2.47 | 66.6 | 12.7 | |
| BgtE-20010 | - | 1 | Candidate effector | E001 | -1.04 | 515.9 | 263.1 | yes |
| Bgt-70087 | - | 1,7[i] | Candidate effector | E001 | -0.68 | 224.5 | 147.9 | |
| Bgt-70088 | - | - | Candidate effector | E001 | -1.04 | 515.9 | 263.1 | |
| BgtE-20010b | - | - | Candidate effector | E001 | -1.04 | 515.9 | 263.1 | |

[a]Signal peptide prediction according to SignalP5.0
[b]Number of amino acid polymorphisms between the parental isolates 96224 and THUN-12
[c]Classification of gene into candidate effector (see (2)). For non-effector genes, the protein was blasted against the NCBI database, putative function is indicated
[d]Effector gene family definition according to (2). NC designates genes that were newly annotated compared to the annotation presented in (2) but like effectors contain a signal peptide and show no homology outside the genus.
[e]Differential gene expression analysis comparing expression levels in *B.g. tritici* 96224 on the susceptible wheat cultivar 'Chinese Spring' at 2dpi with *B.g. triticale* THUN-12 on the susceptible triticale cultivar 'Timbo'. Expression differences are indicated as logFC. logFC >1.5 was considered significant. Genes with missing logFC values are not expressed.
[f]Average gene expression of three biological replicates in isolate 96224 at 2dpi on the susceptible wheat cultivar 'Chinese Spring'. Expression values are indicated as rpkm (Reads Per Kilobase of transcript, per Million mapped reads).
[g]Average gene expression of three biological replicates in isolate THUN-12 at 2dpi on the susceptible triticale cultivar 'Timbo'. Expression values are indicated as rpkm (Reads Per Kilobase of transcript, per Million mapped reads).
[h]Tested in co-expression assays with *Pm17*-HA in *Nicotiana benthamiana.*
[i]Gene is present in two copies in isolate THUN-12.

**Dataset S1 (separate file).** Sequence of effector genes used for expression in *N. benthamiana*

**Dataset S2 (separate file).** List of primers used in this study

**Dataset S2 (separate file).** Information about *B.graminis* isolates used in this study
